# Stochasticity in mammalian cell growth rates drives cell-to-cell variability independently of cell size and divisions

**DOI:** 10.1101/2025.06.18.659700

**Authors:** Ethan Levien, Joon Ho Kang, Kuheli Biswas, Scott R. Manalis, Ariel Amir, Teemu P. Miettinen

## Abstract

Cell growth rates exhibit cell-intrinsic cell-to-cell variability, which influences cell fitness and size homeostasis from bacteria to cancer. Whether this variability arises from noise in cell growth or cell division processes, or originates from cell-size-dependent growth rates, remains unclear. To separate these potential sources of growth variability, single-cell growth rates need to be examined across different timescales. Here, we study cell-intrinsic size and growth regulation by tracking lymphocytic leukemia cell mass accumulation with high precision and minute-scale temporal resolution along long ancestral lineages. We first show that cell-size-dependent growth regulation and asymmetric division of cell size do not explain cell-to-cell growth variability. We then isolate growth fluctuations from overlapping cell-cycle-dependent growth using a Gaussian process regression analysis. We find that these growth fluctuations drive cell-to-cell growth variability within ancestral lineages despite being independent of cell divisions, cell cycle, and cell size. Overall, our results indicate that cell-intrinsic long-term patterns in cell growth are a byproduct of short-term growth fluctuations.

## Main text

All cell populations exhibit cell-to-cell variability in growth [1, 2, 3, 4, 5, 6, 7, 8] and in the underlying growth driving processes, such as cell signaling, transcription, and translation [1, 4, 9, 10, 11, 12, 13, 14]. Understanding the variability between individual cells can elucidate many aspects of physiology and population dynamics. For example, information on cell-to-cell variability can help discriminate between different models of cell growth, gene expression, and size-control [9, 15, 16, 17, 18]. In bacteria and many cancers, cell-to-cell growth variability can also impact drug treatment responses, as non-genetic low-growth states, such as “*persister* “ cells [19], are often more resilient to drugs [8, 17, 20, 21]. Thus, understanding how cell-to-cell growth variability arises even between genetically identical cells is of clear importance.

A key requirement for studying cell-to-cell growth variability is precise growth measurements that enable growth rate monitoring on both short (within a cell cycle) and long (between generations) timescales. Knowledge about measurement precision is also required, as growth fluctuations could falsely arise from poor measurement precision or stability. Tracking single-cell volume growth rates over long timescales has previously been achieved in single-celled organisms using ‘mother machines’ [1, 2]. However, this is not the case for cell mass growth, and cell volume can fluctuate separately from mass [22]. Importantly, there are no long-term (≥4 generations) single-cell growth rate tracking data for mammalian cells, as existing mother machines are not suitable for mammalian cells. Here, we overcome these technical limitations by utilizing suspended microchannel resonators (SMR) to track the mass accumulation rate of leukemia cells across many generations. The SMR is a non-invasive buoyant mass sensor with ∼0.1% mass measurement precision [23, 24, 25]. Buoyant mass is analogous to dry mass (near-perfectly correlated in interphase) [26] and, from here on, we refer to it simply as ‘mass’.

The SMR enabled us to monitor single-cell mass accumulation over timescales ranging from minutes to a week (Fig 1A), allowing us to determine cell-to-cell growth variability (Figs 1B). Using these cell mass (*M*) dynamics, we can obtain instantaneous growth rates (*λ*), and the averaged growth rates within each cell cycle 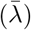. These are defined as [27]

**Figure 1.**
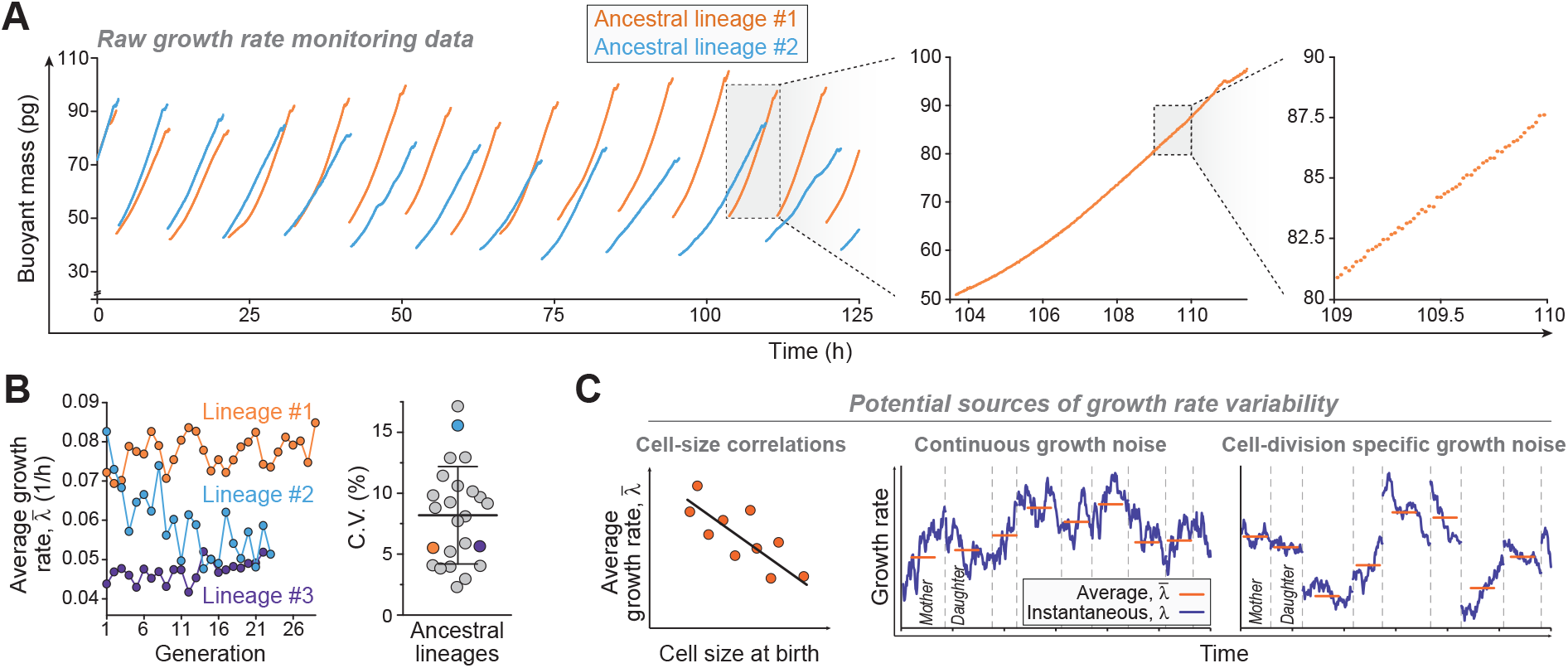
Cell-to-cell growth rate variability in ancestral lineages. (**A**) Two example sections of buoyant mass traces in ancestral L1210 cell lineages collected using the SMR. Buoyant mass of an isolated cell was measured approximately every minute under steady environmental conditions. At each division one daughter cell is randomly discarded. Zoom-ins to a single cell cycle and to a 1-hour section are shown on the right. (**B**) *Left*, average cell growth rates 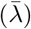 across the three longest lineages. *Right*, coefficient of variability (C.V.) of 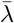 in all lineages (N=24 independent lineages). Bar and whiskers depict mean±SD. (**C**) A schematic of the potential sources of cell-to-cell growth rate variability. Cell-to-cell growth rate variability may arise from cell-size-dependent growth. Alternatively, this variability could also arise from noise in cell growth. Such noise may be continuous, or specific to cell divisions (vertical dashed lines indicate cell divisions). To understand where cell-to-cell growth rate variability arises from, we must resolve the ‘structure’ of cell growth.

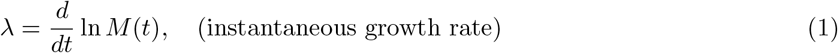

and

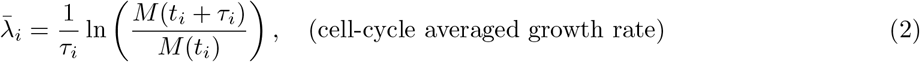

respectively, where *t*_*i*_ is the time at which the *i*th cell was born, *τ*_*i*_ is its cell cycle duration.

To understand cell-to-cell growth variability, we aim to identify deviations from strictly exponential growth in Equation 1. These deviations can arise from cell-size-dependent growth rates, which can reflect size-control (Fig 1C) [28]. Alternatively, deviations from exponential growth can also arise from noise in the biochemistry driving cell growth, resulting in continuous noise in growth rates (Fig 1C) [4, 13, 29]. If this noise is sufficiently large, it may influence cell-to-cell growth variability. However, noise in the underlying biochemistry of the cell can also result in unequal partitioning of cellular components during cell division [13, 29, 30, 31, 32], which could impact cell-to-cell growth variability by imposing growth rate changes specifically at cell divisions (Fig 1C). Therefore, a key question is whether continuous noise, *i*.*e*., fluctuations, in growth rates can be detected and whether they are “blind” to cell divisions.

Here, to disentangle these potential sources of cell-to-cell growth variability, we carry out a comprehensive characterization of the structure of mammalian cell growth (*i*.*e*., overlapping deterministic patterns, degree of randomness, timescale of growth fluctuations), which is currently largely undefined. Using high-precision, single-cell growth rate measurements of leukemia cells, we first show that most cell-to-cell growth variability is not explained by cell-size-dependent regulation of 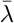 or division asymmetries. Next, we utilize Gaussian process (GP) regression analysis to infer and isolate growth fluctuations that are independent of longer timescale growth regulation as well as measurement fluctuations. These fluctuations reveal that continuous noise in growth, rather than cell division-specific changes, explains the majority of cell-to-cell growth variability in an ancestral cell lineage.

## Results

### Monitoring cell growth across ancestral lineages

Using the SMR, we collected a large dataset of mouse lymphocytic leukemia L1210 cell growth rates. Our data contains 24 lineages (each lineage is an independent experiment) with a total of 235 full cell cycles and 2600 h of data (Fig S3A, Table S1). Optimization of system stability allowed us to monitor growth in ancestral lineages ranging from 3 to 29 full generations in length with cell mass measurements every ∼1 minute. The SMR is not capable of tracking the growth of both sister cells following cell division, and one of the daughter cells is randomly discarded following every division. This data represents, to the best of our knowledge, the longest instantaneous growth rate monitoring data for mammalian cells to date. Furthermore, because our growth rate measurements reflect mass accumulation, our results are not sensitive to volume fluctuations reported for many mammalian cells [33, 34].

We first characterized the overall growth behavior of the cells. The time that cells spent in the SMR did not correlate with 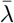 (Fig S3B). The cell cycle durations (*τ* = 10.2±2.0 h, mean±SD) were comparable to independent studies with L1210 cells [23, 35], and the amount of variability in *τ* was comparable to previous work where all data were collected at the same time (Fig S3C) [36]. All L1210 cells in bulk culture were positive for the proliferation marker Ki-67 (Fig S3D) and all cells measured with SMR were growing. A cell in the SMR is in isolation from other cells and exposed to fresh media with every mass measurement, thereby excluding growth regulation due to autocrine and paracrine signals [37, 38]. To focus our analyses on steady-state growth, the first (partial) cell cycle of each lineage was removed. We also examined the technical stability of our SMR setup by repeatedly measuring the mass of polystyrene beads. We did not observe systematic fluctuations in bead mass (Fig S3E), and our measurement calibration drifted only 0.07% during a typical cell cycle. Overall, our ancestral lineage dataset reflects cancer cells growing under steady, high-growth conditions, where all cells actively proliferate, and growth behaviors can be attributed to cell-intrinsic sources.

When examining the generic characteristics of cell-to-cell growth variability, we observed that cells within each lineage had a similar 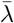, although lineages differed in their average growth rate by *∼* 11% (Fig. 1B, Fig S3A). Experiments monitoring the proliferation rate of distinct L1210 cell lineages have also reported similar observations, where each cell lineage proliferates at its lineage-specific rate [8, 36]. On average, the cell-to-cell growth variability within each lineages was ∼8% (Fig. 1B). Here, we focus on explaining this within-lineage growth variability.

### Connections between size-control and cell-to-cell growth rate variability

Cell size-control is often studied within an autoregressive framework [28, 39], where the cell size-control strategy is defined by a parameter *α*. This relates the cell’s birth mass (*M*_0_) to the daughter cell’s birth mass (*M*_*f*_):

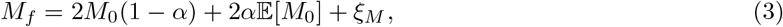

where 𝔼 [*M*_0_] is the average initial mass, or equivalently the mass added during the cell cycle, and *ξ*_*M*_ is a Gaussian random variable. Values of *α* range from 0 to 2 with lower *α* values indicating a weaker size homeostasis [39]. Many cell types are reported to follow the adder model of size-control (*α* ≈ 0.5) [5, 6, 40, 41, 42, 43, 44, 45, 46], where cells grow an approximately constant amount in each cell cycle.

Using our new dataset, we estimated *α* (Eq. 3) in each lineage. On average, our data is consistent with an adder size-control strategy (*α* = 0.55±0.06, mean±SE, N=21, 3 shortest lineages excluded). This *α* can be achieved by regulating *τ* and/or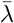. When examining these size-dependencies, we observed that almost all individual lineages have a negative correlation between cell birth mass and *τ* (Pearson *R*^2^ = 0.25±0.07, mean±SE, Fig 2A), but not between between cell birth mass and 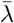 (Pearson *R*^2^ = 0.10±0.04, mean±SE,

**Figure 2.**
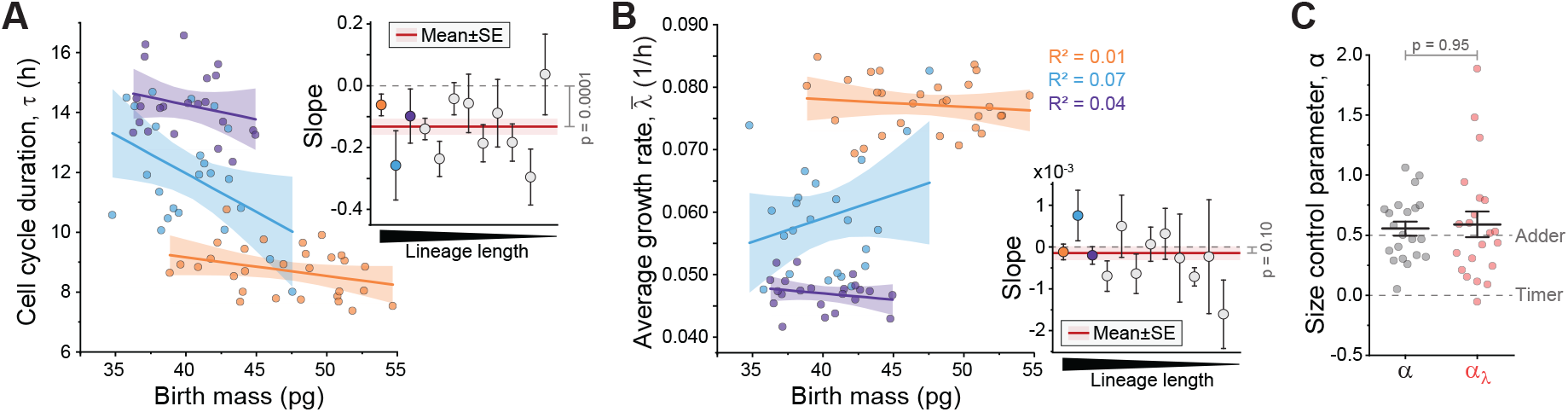
Size-control does not explain cell-to-cell growth variability. (**A-B**) Correlation between cell birth mass and *τ* (A) or 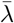 (B) in the three longest lineages. Each inset displays the slope of the correlation for all lineages with ≥8 cells (N=12 independent lineages). Inset red line and shaded area depict mean±SE (data weighted by lineage length), p-value depicts one sample t-test in comparison to 0. (**C**) Cell size-control parameter, *α* when analyzed normally (*α*, grey) and when conditioning the data on 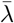 to only considering the contribution of cell cycle duration variability on *α* (*α*_*λ*_, red). Dots depict independent lineages (N=24), bar and whiskers depict mean±SE, p-values obtained by Welch’s t-test.

Fig 2B). To quantify the relative contribution of 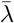 to the overall size-control, we followed the approach of Cadart et al. [5] and used the scaled regression coefficient of initial size on average growth rate as a measure of the role of growth in size-control. The regression coefficient is given by

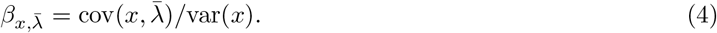

To obtain a non-dimensional parameter, we scale by the average generation time to obtain 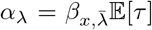. Computing *α*_*λ*_ revealed that conditioning the data on the growth rate had no impact on the strength of size-control (Fig 2C). Thus, in L1210 cells, cell-size homeostasis can be maintained by the regulation of *τ* alone, and the variability in cell-to-cell growth rates cannot be explained by size-control.

### Cell-to-cell growth variability is independent of cell division symmetry

We next examined if asymmetric cell mass divisions, where the mass of the newborn daughter cell is not perfectly 1/2 of the mass of the dividing mother cell, could explain the cell-to-cell growth variability. In each lineage,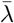 typically changed ∼7% from mothers to daughters (Fig 3A), and cell mass divisions displayed, on average, ∼ 3% deviation from perfectly symmetrical divisions (Fig 3B). When we correlated the division asymmetry with the change in 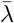 in each mother-daughter pair, we did not observe any correlation (Fig 3C). In addition, the lineages displayed little memory of 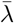 and the autocorrelation function of 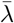 decayed to zero in two generations (Fig 3D). Thus, the variability in 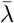 is not explained by systematic patterns (memory) in 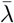 or by asymmetric cell mass divisions. We note that this degree of division asymmetry is very low, and previous results have suggested over 2-fold higher asymmetry in cell mass divisions across cell lines [47].

**Figure 3.**
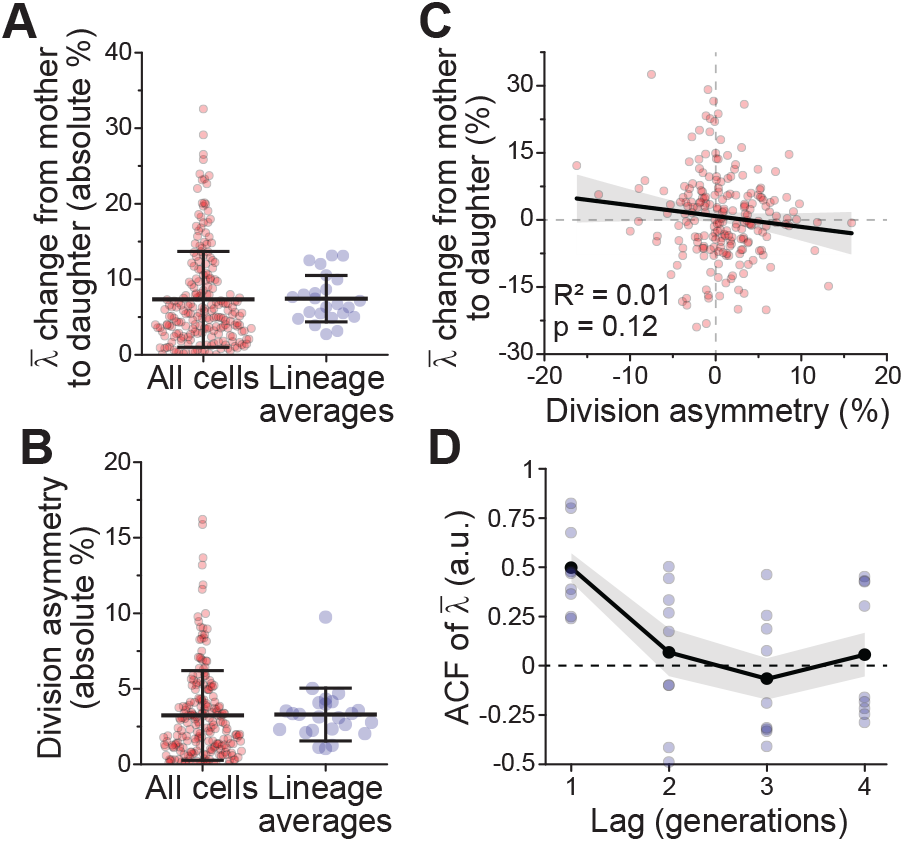
Cell-to-cell growth variability is independent of cell mass division errors. (**A**,**B**) Absolute % change in 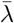 from mother to daughter (A), and absolute % cell mass division asymmetry (i.e. deviation from perfectly symmetrical division of cell mass) (B). Data is shown separately for all cells (red, n=211 mother-daughter pairs) and the averages of each lineage (blue, N=24 independent lineages). Bar and whiskers represents mean±SD. (**C**) Correlation between cell mass division asymmetry and the change in 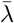 from mother to daughter. Each dot represents a mother-daughter pair (n=221). Black line and shaded area depict linear fit and 95% confidence intervals. Pearson correlation and p-value (ANOVA) are also displayed. (**D**) Autocorrelation of 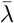 in the 10 longest lineages. Black line and shaded areas represent mean±SEM.

### Isolation of growth fluctuations

Having excluded size-control and mass division asymmetry as potential explanations for cell-to-cell growth variability, we moved to examine the structure of growth within the cell-cycle in more detail. As discovered in a previous study [48], we observed that each lineage had a cell-cycle-dependent growth trend that was dependent on the relative age in the cell cycle, not the absolute time since cell birth (Fig S4). This results in *λ* being cell-size and cell-cycle-dependent. To isolate fluctuations in *λ* around the cell-cycle-dependent growth trend, we used a Gaussian process (GP) regression analysis. GP is a a Bayesian, kernel-based method which can be used to simultaneously smooth and decompose a function *f*, as shown before using different types of growth measurements [49, 50]. The GP is a based on a series expansion of the form *f* (*t*) = ∑ _*i≥*0_ *β*_*i*_*φ*_*i*_(*t*) where {*φ*_*i*_(*t*)} represent basis functions. Instead of specifying these functions directly, one places Gaussian priors on *β*_*i*_. The posterior distribution 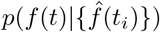 (the distribution of the function values *f* conditioned on the data) can then be obtained analytically by marginalizing over the *β*_*i*_. This distribution is determined only by the covariance function

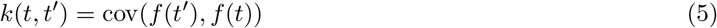

which is selected based on assumptions about the signal. The sum and derivatives of a Gaussian process are also Gaussian processes, and the process inherits the smoothness properties of the kernel [51]. Consequently, we can use the isolated mass terms of GP to infer *λ*. Unlike other smoothing procedures, such as spline smoothing or Savitzky-Golay filters, in GP, the error in the mass measurements does not propagate, leading to spurious correlations in the inferred *λ*.

Our specific GP pipeline (see ***SI Appendix, section 2***) is as follows (Fig 4A): First, we paste together the log masses to obtain a continuous signal:

**Figure 4.**
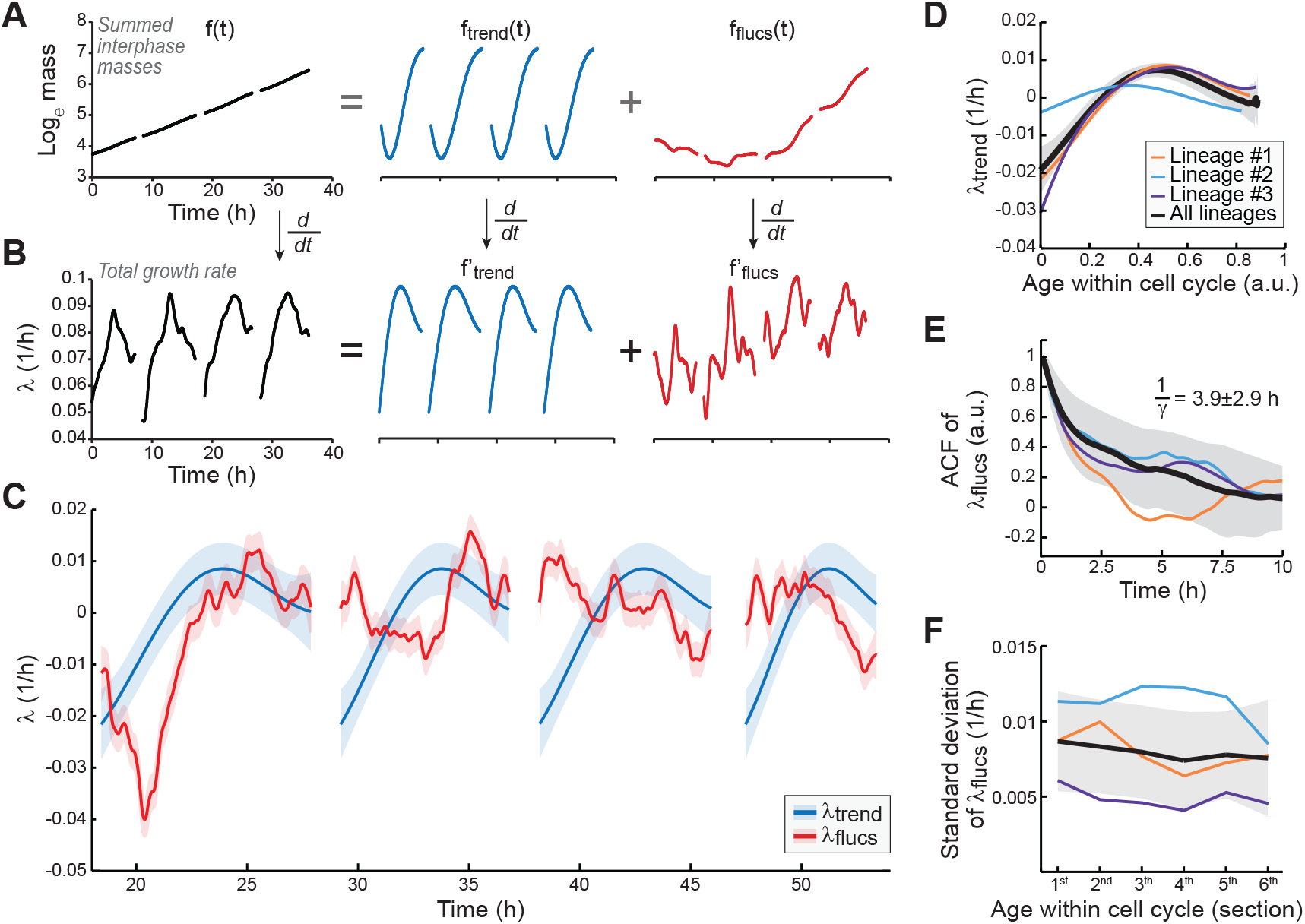
Growth rate fluctuations are independent of the cell-cycle. (**A-B**) Analysis workflow: (A) Cell mass traces are plotted on a logarithmic scale; mitoses are removed and the mass traces are summed. Gaussian process is used to decompose the summed masses into discrete terms that reflect the cell-cycle-dependent mass trend and mass fluctuations. (B) The isolated mass terms are converted to instantaneous growth terms. (**C**) An example of the isolated instantaneous growth terms over four full cell cycles in lineage #1. The gaps in data are cell divisions. Solid lines and shaded areas display the mean ± SD of each posterior distribution. The data is normalized to mean-zero. (**D**) The cell-cycle-dependent *λ*. (**E**) Autocorrelations of the *λ* fluctuations within each cell cycle. (**F**) Magnitude (standard deviation) of the *λ* fluctuations in different cell cycle sections (i.e. at different cell age). In panels D-F, colored lines depict the three longest lineages and black line with shaded areas depict all lineages (mean ± SD, N=24 independent lineages).

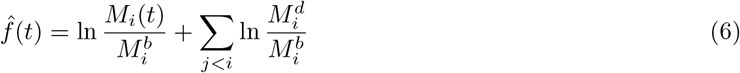

where 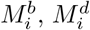 nd *M*_*i*_(*t*) are the masses of the *i*th cell in a lineage at cell birth, cell division and (absolute) time *t*, respectively. Due to the complex growth dynamics in mitosis [25, 26], we excluded mitoses from our analysis. Our GP decomposes the log summed mass measurements as

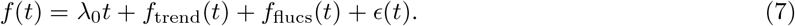

where *c* is a delta correlated Gaussian noise term.

This GP-based approach enabled us to isolate cell mass and *λ* behaviors (Fig 4A-B). The posterior distributions of the cell-cycle-dependent 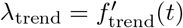 (*t*) and the fluctuations 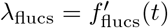 were narrow enough to separate changes in these growth behaviors on a sub-hour timescale (Fig 4C). Before analyzing these growth behaviors in more detail, we verified that the cell mass and *λ* fluctuations reflect changes in cell mass rather than measurement error. We found that consecutive residuals of our GP analysis were not correlated (Fig S5A, *R*^2^=0.0046). The degree of these correlations was comparable to the degree of correlations observed in polystyrene bead measurements (Fig S5B), which is the level expected based purely on measurement noise. However, we note that lineage # 2 displayed higher correlations between consecutive residuals than other lineages and data from this lineage may be skewed. We also observed that the GP-isolated mass fluctuations in cells are orders of magnitude higher than those observed in polystyrene beads (Fig S5C-D) and that the cell mass fluctuations took place on several orders of magnitude longer timescales than the bead mass fluctuations (Fig S5E). Thus, the GP-isolated cell mass and *λ* fluctuations do not reflect measurement artifacts but rather ‘true’ growth behaviors.

### Growth fluctuations are independent of the cell-cycle and cell-size

We examined the GP-isolated growth terms to understand their impact on *λ*. The cell-cycle-dependent *λ*_trend_ was typically maximized in the middle of each cell cycle (at age 0.5±0.1, mean±SD, Fig 4D). From cell birth to the point of maximum growth, the cell-cycle influenced *λ* by 0.028±0.008 1/h (mean±SD), which corresponds to 38±14% (mean±SD) of total *λ*. Similar conclusion was reached when growth rates were analyzed using a simple fitting approach (Fig S6). The influence that the *λ*_trend_ and the *λ*_flucs_ had on the total *λ* were comparable. However, as our GP analysis assumes every cell in a given lineage displays the same *λ*_trend_ (after normalization for cell age), essentially all cell-to-cell growth variability is included in the *λ*_flucs_. This allows us to focus on the *λ*_flucs_ as we aim to understand cell-to-cell growth variability.

When examining the autocorrelation of *λ*_flucs_, we did not observe any periodicity (Fig 4E). This is in contrast to a recent study, which also examined mass growth in L1210 cells, proposing that cell growth undergoes oscillations [52]. The *λ*_flucs_ displayed a relaxation timescale (1*/γ*) of ∼4 hours whether analyzed from the autocorrelation function or from the variance in the magnitude of the fluctuations (***SI Appendix, section 1)***. These relaxation timescales are shorter than *τ* and thereby consistent with 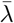 having little memory over generations.

Next, we looked for cell-cycle-dependency in *λ*_flucs_. We did not observe any difference in the magnitude (standard deviation) of *λ*_flucs_ between the cells of different age in the cell-cycle. Thus, the *λ*_flucs_ are independent of the cell-cycle state (and approximate cell-size) (Fig 4F), even though the total growth rate of cells is not (Fig 4B). Previous studies in bacteria and mammalian cells have suggested that cell volume growth variability is higher in small cells at the beginning of the cell cycle than in large cells later in the cell-cycle [27, 33]. As we do not observe this, cell-cycle-dependent cell mass behavior might be distinct from cell-cycle-dependent volume behavior. To further analyze if the *λ*_flucs_ displayed cell-size and/or growth-dependency, we performed regressions on the magnitude of the *λ*_flucs_ with cell’s birth mass, division mass, the mass added in a cell-cycle or *τ* (Fig S7). We did not observe any statistically significant correlations. Overall, these results suggest that the *λ*_flucs_ do not reflect cell-size or cell-cycle -dependent growth regulation, nor do the *λ*_flucs_ display any clear patterns.

We have also used our data to analyze cell-to-cell variability in *λ* on different timescales. We detail this analysis and its potential impact in the ***SI Appendix, section 3***.

### Cell-to-cell growth variability is predicted by cell-division-independent growth fluctuations

Our isolation of *λ*_flucs_ provides us with an opportunity to examine if cell growth rates are sensitive to cell division events. We compared the observed cell-to-cell variability in 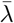 to the prediction of a simple null model in which instantaneous growth rate fluctuations are causally decoupled from the cell-cycle. Specifically, we modeled *λ*_flucs_(*t*) as an Ornstein–Uhlenbeck (OU) process – essentially a continuous time mean-reverting random walk [53] – which was motivated by the exponential decay in *λ*_flucs_ autocorrelations and the lack of correlations between *λ*_flucs_ and cell-size or age (Fig 4E, Fig S7). An OU process can be defined mathematically by the stochastic differential equation

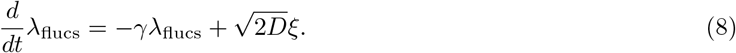

Here *ξ* is a white noise term, *D* is the diffusion coefficient and *γ* is the relaxation rate. At cell-division, we assumed that *λ*_flucs_ is perturbed by a Normal random variable *z*.

This model interpolates between two simple limits: As the growth noise introduced at cell division (*σ*_*z*_) → 0 we are left with the standard OU process which is “blind” to cell divisions (continuous noise). As *D* → 0 we obtain a model where growth variation is tied to the cell division events. The contribution of the continuous growth fluctuations and division-specific growth noise can be identified by looking at the time averaged growth rate,

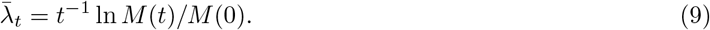

where *t* = 0 is when cell division occurs. The variance of 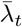 can be decomposed as

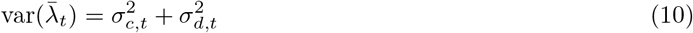

where *σ*_*c*_ and *σ*_*d*_ respectively represent the variation from the OU and division noise. When the characteristic relaxation time-scale (1*/γ*) of this process is much less than the cell age (the absolute time since cell birth), these are given by (see ***SI Appendix, section 1***)

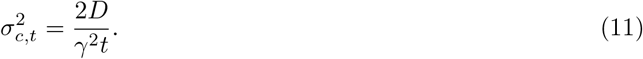

and

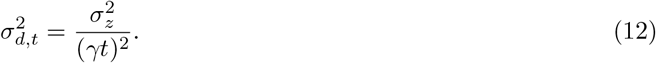

We compared these expressions to the experimental data by examining the scaling between 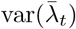 and cell age (*t*) (Fig 5A). The data favor the former model where scaling between 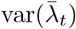 and cell age (*t*) has an exponent of -1 (Fig 5A, black dashed lines), as growth variation is blind to cell divisions. This conclusion persisted whether *λ*_flucs_ was isolated with GP or analyzed directly from raw data (Fig 5A, red solid lines). We note, however, that we observed scaling exponents lower than those predicted by pure OU noise, especially when analyzing the GP isolated *λ*_flucs_.

**Figure 5.**
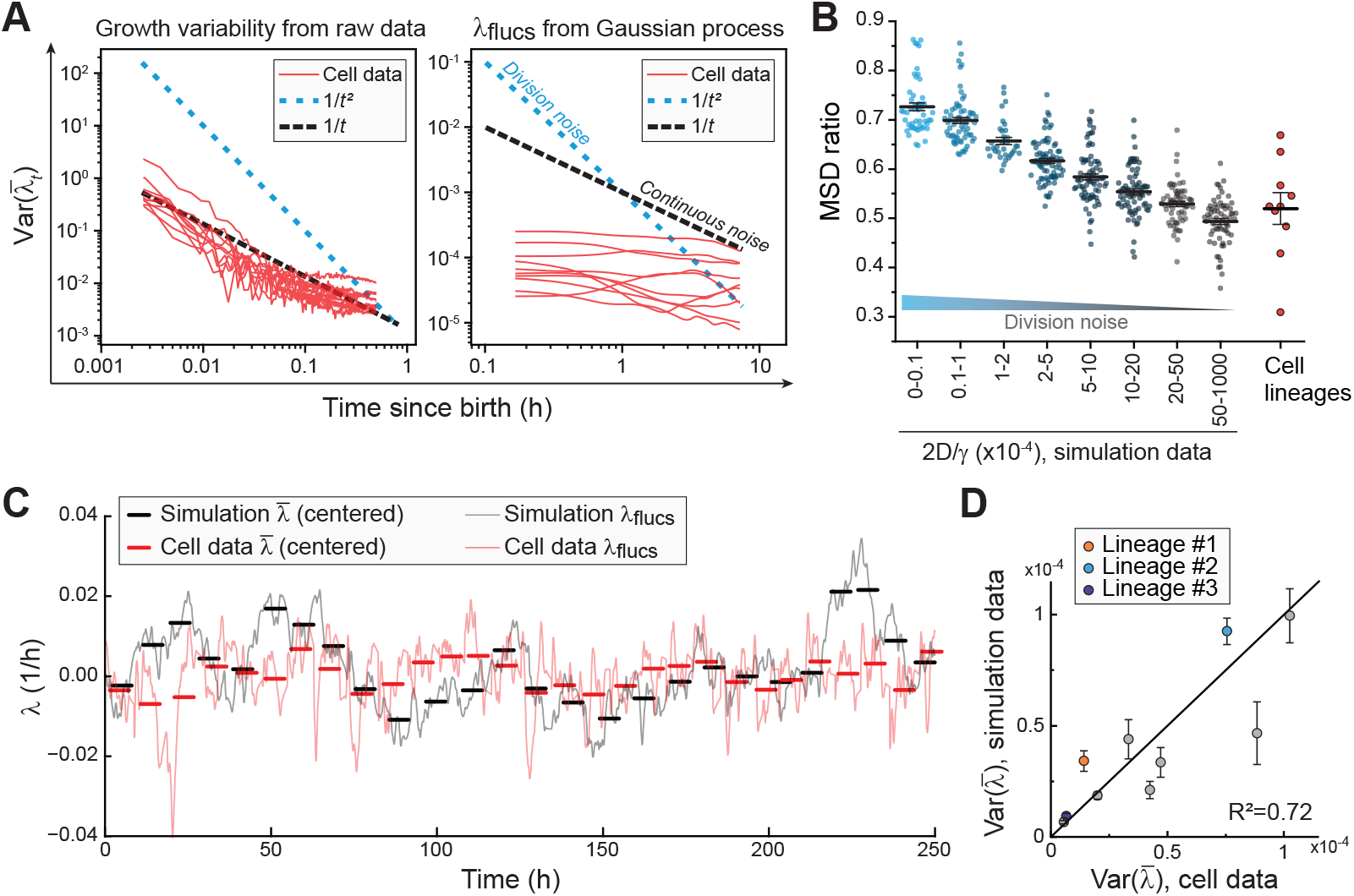
Cell-to-cell growth variability within ancestral lineages arises predominantly from continuous growth fluctuations (**A**) Scaling of 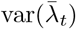 with age (*t*) as observed in raw data (left, N=24) and GP-isolated *λ*_flucs_ (right, N=10 longest lineages). Each red lines depicts a separate lineage, while dotted lines depict model predictions. (**B**) MSD ratio in simulated datasets (teal/grey) and in cell data (red). Simulated data was pooled based on the ratio between *D* and *γ*, so that high values indicate that continuous cell growth noise dominates over cell division noise. Line and error bars represent mean±SEM. Each dot depicts a separate lineage (for cell data, N=10 longest lineages). (**C**) An example of the *λ*_flucs_ in lineage #1 (continuous red line) and in a corresponding OU model simulated data, where growth is ‘blind’ to cell divisions (continuous black line). Thick horizontal bars indicate 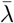 for each cell. Data is normalized to mean-zero. (**D**) Correlation of cell-to-cell variability in 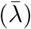 between the cell data and the simulated data with continuous OU noise. Black line indicates x=y. Each dot depicts a separate lineage (N=10 longest lineages), and error bars depict SEM.

We then generated simulated growth data with graded levels of continuous and division noise, and analyzed them using the GP regression to isolate fluctuations in *λ*. We interpolated between the two limits by fixing the cell-cycle averaged growth rate 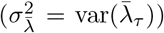 and varying 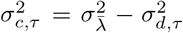. Here *τ* is the generation time. Our simulated data sets were subject to the same GP smoothing as the experimental data.

To compare the simulated data from these models with the experimental L1210 cell data, we calculated the mean squared displacement (*MSD*) of *λ*_flucs_ within each cell, *MSD*_within_, by calculating the sqaured difference in *λ*_flucs_ values across time lags restricted to individual cell cycles. Separately, we computed *MSD*_between_ by calculating the squared difference in *λ*_flucs_ values across equivalent time lags between different cells. The MSD ratio, *MSD*_between_/(*MSD*_within_+*MSD*_between_), separates the simulated datasets using different combinations of *D* and *σ*_*z*_ (Fig 5B). Thus, while the GP can suppress the fluctuations on the timescale of a cell-cycle (Fig 5A), it preserves the growth rate jumps at cell division, allowing us to separate the contributions of continuous and division noise. Using the MSD ratio, we observed that the cell data is indistinguishable from the limiting case simulations where *σ*_*z*_ = 0 (Fig 5B). Thus, growth noise is not introduced at cell divisions but rather continuously throughout the cell cycle, akin to an OU process.

Finally, to test whether a simple OU model predicts the cell-to-cell growth variability within cell lineages, we fitted the parameters in the OU model using a loss function that depends only on fluctuations within the cell cycle. In particular, we fit *γ* and *D* by matching the variance of 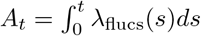 between the model and the data, since *A*_*t*_ has smaller error bars than *λ*_flucs_. Using the fit parameters for each lineage, we simulated lineages with a similar number of cells and time resolution as our experimental data (Fig 5C). We then compared the predicted cell-to-cell variability in 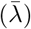 in the simulated lineages with the experimental data, and we observed a strong correlation (Pearson *R*^2^ = 0.72, Fig 5D). We conclude that cell-to-cell variability in 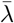 can be largely explained by the instantaneous growth rate fluctuations, which are independent of cell divisions and size.

## Discussion

Here, we have elucidated the structure of cell growth by monitoring single leukemia cell mass accumulation rate across ancestral lineages. The high precision and temporal resolution of our data has allowed us to isolate overlapping but independent growth behaviors. We have verified that the growth behaviors we observe cannot be explained by technical measurement fluctuations, and our experiments are carried out under steady external conditions using multiple independent setups. Thus, our results reflect predominantly cell-intrinsic behaviors in biomass growth. However, our work is not without limitations, and we discuss these in the ***SI Appendix, section 4***.

We find that cell-to-cell growth variability within ancestral lineages can be explained by a two parameter model where the fluctuations in instantaneous growth rates are independent of cell divisions and cell size. In contrast, size-dependent growth rates, asymmetric divisions of cell mass, and growth rate perturbations at cell divisions do not explain the cell-to-cell variability in growth. Notably, modeling studies on imperfect cell divisions have suggested that partitioning errors introduce significant cell-to-cell variability [4, 13, 29, 30], and it would be reasonable to assume that this extends to cell growth rates. Indeed, recent studies in bacteria [27, 54] and mammalian cells [33] have revealed that cell volume growth fluctuation are larger in newborn cells than later in the cell cycle, suggesting that cell divisions may introduce growth noise. Our results on cell mass growth argue against this conclusion in the L1210 leukemia cells. The separate regulation of cell volume and cell mass growth may explain these discrepancies [22], but we also note that our cell line displays less cell mass division asymmetry than many other mammalian cell lines [47].

More broadly, our work shows that long-term cell-to-cell growth variability is actually a byproduct of continuous short-timescale stochasticity in growth. In a simplistic and hypothetical model, where growth rates are set by a single growth limiting-molecule, our results suggest that this growth limiting-molecule is either very high in abundance or that cells possess mechanisms that limit partitioning errors of the molecule. Cells have mechanisms to promote symmetric partitioning of certain cell cycle regulators [55, 56] and cell organelles [57], making it plausible that also key growth regulators undergo similar partitioning.

## Materials and Methods

### Cell culture conditions

The L1210 cells were obtained from ATCC (#CCL-219) and validated negative for mycoplasma. All SMR experiments and the maintenance of cell cultures were carried out in RPMI 1640 medium (Invitrogen, #11875093), supplemented with 10% heat inactivated FBS (Sigma-Aldrich, #F4135, Lot#13C519), 10 mM HEPES (Invitrogen, #15630080), 1 mM sodium pyruvate (Invitrogen, #11360070) and 1x Antibiotic-Antimycotic (Invitrogen, #15240112). All experiments were started when cells were at non-confluent and exponentially growing concentration (100.000 to 500.000 cells/ml).

Cell cycle and proliferation status were examined by fixing cells with 4% PFA for 10 min, permeabilizing cells with 0.25% Triton X-100 for 10 min, washing and blocking the cells with PBS supplemented with 5% BSA for 15 min. The cells were labeled for Ki-67 with 1:250 diluted Alexa Fluor 488 conjugated anti-Ki-67 rabbit monoclonal antibody (Cell Signaling Technologies, #11882) o/n, washed with PBS supplemented with 5% BSA for 15 min, then labeled for DNA with FxCycle PI/RNase staining solution (Invitrogen, #F10797) for 30 min. The cells were analyzed using a flow cytometer (LSR II HTS, BD Biosciences, 488 nm and 561 nm excitation lasers, 530/30 and 610/20 emission filters).

### SMR operation

The SMR chip was mounted on a metal holder connected to a 37°C water bath to maintain constant temperature. The cells were loaded into the SMR from vials pressurized with 5% CO2 and 21% O2. The fluidic pressure system was set to replenishes fresh media into SMR with every cell measurement, approximately every 1 min, thereby maintaining steady conditions throughout the experiment. Full details of the SMR setup, operation and frequency data analysis can be found in [48, 25, 35].

The following changes were implemented to increase the long-term stability of the single-cell hydrodynamic trap: 1) Fresh media in SMR was replenished directly from 20 ml glass Wheaton vials to minimize the fluid-height driven pressure difference during the trap. 2) The glass Wheaton vials were place on micrometers to tune fluid height daily throughout the experiments. 3) The glass Wheaton vials were not heated as heating accelerates water evaporation from the media. 4) Immediately after a cell was loaded into the SMR, we flushed other cells out of the SMR and the tubing using a flow rate of more than 10 nl/s and taping of the tubing for approximately 2 min. 5) After the initial cell loading step, the SMR was kept in the dark to remove any potential phototoxicity.

Across the study, three independent SMR setup were used to collect the data and no systematic differences were observed between the results of each setup.

### System calibration

The SMR systems were calibrated using two approaches [48, 25, 35]. First, the SMR was loaded with sodium chloride solutions (0, 2, 6, 10 and 16% w/v) for which fluid densities are known. The SMR resonant frequency was measured at RT using open-loop setting to generate a baseline solution density calibration curve. Second, the change in SMR resonant frequency to an object of known buoyant mass was calibrated to derive a mass calibration factor (Hz/pg). 10 µm diameter polystyrene beads of density 1.05g/cm^3^, suspended in DI water or PBS, were used as calibration particles.

## Autocorrelations

Autocorrelation coefficients were obtained by using the following equation.

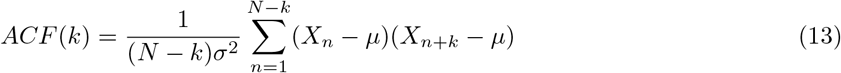

Where *X*_*n*_ is the data (i.e. either *λ* or 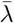). *N* and *σ* are the total number of data points and the standard deviation of *X*_*n*_, respectively. For plotting the ACF of *λ* as a function of time, we normalized the discrete time step between adjacent data points of the above equation by the median value of the time difference between the adjacent data points in our measurement or simulation (*t* = *kx*_*cal*_, where *x*_*cal*_ = *median*(Δ*t*)).

### Mean Squared Deviation (MSD) analysis

The mean squared deviation (MSD) of the fluctuations in *λ* for a time lag *τ* is defined as below,

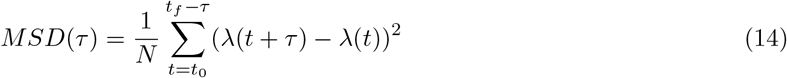

where *N* is the total number of data summed between the first time point *t*_0_ and the last time point *t*_*f*_ subtracted by the time lag *τ*. MSD was computed for each lineage.

The *MSD*_*within*_ of the fluctuations in *λ*_*i*_ was calculated by only computing the MSD values within each cell *i* in the lineage. In other words,

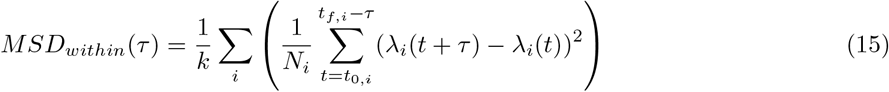

where *k* is the total number of cells within the lineage, *N*_*i*_, *t*_0,*i*_ and *t*_*f*,*i*_ are the total number of data summed, the first time point and the last time point of the data, respectively, for individual cell *i* in the lineage.

The *MSD*_*between*_ of the fluctuations in *λ*_*i*_ and *λ*_*j*_ was calculated by only computing the MSD values across different cells. In other words,

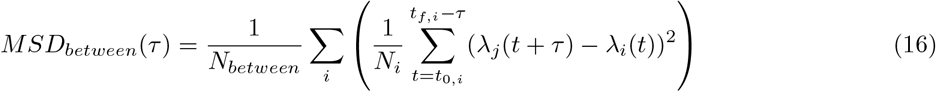

where *j /*= *i*. Note that for each cell *i*, there is at most one other cell *j* for which the deviation *λ*_*j*_(*t*+*τ*) *− λ*_*i*_(*t*) can be calculated. This limitation arises due to the sequential nature of our dataset, where *t*_*f*,*i*_ *< t*_0,*j*_ if *i < j*.

The MSD ratios represent

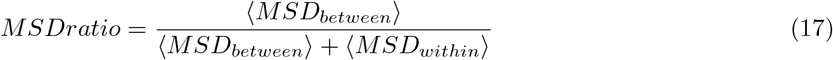

where *< MSD >* indicate values averaged for *τ* between 0 h and 4 h.

## Supporting information

Supplementary Section

## Data and code availability

Data is a combination of our previously published data (7 lineages) [25] and new experimental data (17 lineages), all collected under identical conditions. All single-cell mass traces can be found attached in the online supporting information (Table S1). Codes used for the Gaussian process regression analysis and generation of simulated data can be found at: https://github.com/elevien/L1210

## Acknowledgments

J.H.K received funding and support from the Hyper-Convergence Research Program and Creative-Pioneering Researchers Program via the Institute of Advanced Machines and Design at Seoul National University (SNU), and the National Research Foundation of Korea (RS-2024-00342400) via the Institute of Engineering Research at SNU. A.A. received funding and support from the Clore Center for Biological Physics. S.R.M. received support from Virginia and D.K. Ludwig Fund for Cancer Research and MIT Center for Precision Cancer Medicine. T.P.M. received funding from Wellcome (110275/Z/15/Z). We thank Alice R. Lam for her assistance and constructive feedback.

## Competing interests

S.R.M. is a co-founder of Travera and Affinity Biosensors, which develop techniques relevant to the research presented. The remaining authors declare no competing interests.

## References

[1] D. J. Kiviet, P. Nghe, N. Walker, S. Boulineau, V. Sunderlikova, and S. J. Tans. Stochasticity of metabolism and growth at the single-cell level. Nature, 514(7522):376–9, 2014.

[2] P. Wang, L. Robert, J. Pelletier, W. L. Dang, F. Taddei, A. Wright, and S. Jun. Robust growth of escherichia coli. Curr Biol, 20(12):1099–1103, 2010.

[3] H. Vashistha, M. Kohram, and H. Salman. Non-genetic inheritance restraint of cell-to-cell variation. Elife, 10:e64779, 2021.

[4] P. Thomas, G. Terradot, V. Danos, and A. Y. Weisse. Sources, propagation and consequences of stochasticity in cellular growth. Nat Commun, 9:4528, 2018.

[5] C. Cadart, S. Monnier, J. Grilli, P. J. Saez, N. Srivastava, R. Attia, E. Terriac, B. Baum, M. Cosentino-Lagomarsino, and M. Piel. Size control in mammalian cells involves modulation of both growth rate and cell cycle duration. Nat Commun, 9:3275, 2018.

[6] I. Soifer, L. Robert, and A. Amir. Single-cell analysis of growth in budding yeast and bacteria reveals a common size regulation strategy. Curr Biol, 26(3):356–361, 2016.

[7] M. M. Logsdon, P-Y. Ho, K. Papavinasasundaram, K. Richardson, M. Cokol, C. M. Sassetti, A. Amir, and B. B. Aldridge. A parallel adder coordinates mycobacterial cell-cycle progression and cell-size homeostasis in the context of asymmetric growth and organization. Curr Biol, 27(21):3367–3374, 2017.

[8] A. Seita, H. Nakaoka, R. Okura, and Y. Wakamoto. Intrinsic growth heterogeneity of mouse leukemia cells underlies differential susceptibility to a growth-inhibiting anticancer drug. PLoS One, 16(2):e0236534, 2021.

[9] M. Wallden, D. Fange, E. G. Lundius, O. Baltekin, and J. Elf. The synchronization of replication and division cycles in individual e. coli cells. Cell, 166(3):729–739, 2016.

[10] A. Colman-Lerner, A. Gordon, E. Serra, T. Chin, O. Resnekov, D. Endy, C. G. Pesce, and R. Brent. Regulated cell-to-cell variation in a cell-fate decision system. Nature, 437(7059):699–706, 2005.

[11] M. B. Elowitz, A. J. Levine, E. D. Siggia, and P. S. Swain. Stochastic gene expression in a single cell. Science, 297(5584):1183–6, 2002.

[12] M. Ricicova, M. Hamidi, A. Quiring, A. Niemisto, E. Emberly, and C. L. Hansen. Dissecting genealogy and cell cycle as sources of cell-to-cell variability in mapk signaling using high-throughput lineage tracking. Proc Natl Acad Sci U S A, 110(28):11403–11408, 2013.

[13] A. Schwabe and F. J. Bruggeman. Contributions of cell growth and biochemical reactions to nongenetic variability of cells. Biophys J, 107(2):301–313, 2014.

[14] S. Di Talia, J.M. Skotheim, J.M. Bean, E.D. Siggia, and F.R. Cross. The effects of molecular noise and size control on variability in the budding yeast cell cycle. Nature, 448(7156):947–951, 2007.

[15] B. Cerulus, A. M. New, K. Pougach, and K. J. Verstrepen. Noise and epigenetic inheritance of single-cell division times influence population fitness. Curr Biol, 26(9):1138–47, 2016.

[16] J. Lin and A. Amir. The effects of stochasticity at the single-cell level and cell size control on the population growth. Cell Systems, 5(4):358–367, 2017.

[17] S. Shen, S. Vagner, and C. Robert. Persistent cancer cells: The deadly survivors. Cell, 183(4):860–874, 2020.

[18] A. Amir and N. Q. Balaban. Learning from noise: how observing stochasticity may aid microbiology. Trends in Microbiology, 26(4):376–385, 2018.

[19] E. Kussell, R. Kishony, N. Q. Balaban, and S. Leibler. Bacterial persistence: a model of survival in changing environments. Genetics, 169(4):1807–1814, 2005.

[20] C. Duy, M. Li, M. Teater, C. Meydan, F. E. Garrett-Bakelman, T. C. Lee, C. R. Chin, C. Durmaz, K. C. Kawabata, E. Dhimolea, C. S. Mitsiades, H. Doehner, R. J. D’Andrea, M. W. Becker, E. M. Paietta, C. E. Mason, M. Carroll, and A. M. Melnick. Chemotherapy induces senescence-like resilient cells capable of initiating aml recurrence. Cancer Discovery, 11(6):1542–1561, 2021.

[21] K. A. Fennell, D. Vassiliadis, E. Y. N. Lam, L. G. Martelotto, J. J. Balic, S. Hollizeck, T. S. Weber, T. Semple, Q. Wang, D. C. Miles, L. MacPherson, Y. C. Chan, A. A. Guirguis, L. M. Kats, E. S. Wong, S. J. Dawson, S. H. Naik, and M. A. Dawson. Non-genetic determinants of malignant clonal fitness at single-cell resolution. Nature, 601(7891):125–131, 2022.

[22] J. Lin, J. Min, and A. Amir. Optimal segregation of proteins: Phase transitions and symmetry breaking. Physical Review Letters, 122:068101, Feb 2019.

[23] S. Son, A. Tzur, Y. Weng, P. Jorgensen, J. Kim, M. W. Kirschner, and S. R. Manalis. Direct observation of mammalian cell growth and size regulation. Nat Methods, 9(9):910–2, 2012.

[24] T. P. Burg, M. Godin, S. M. Knudsen, W. Shen, G. Carlson, J. S. Foster, K. Babcock, and S. R. Manalis. Weighing of biomolecules, single cells and single nanoparticles in fluid. Nature, 446(7139):1066–9, 2007.

[25] T. P. Miettinen, J. H. Kang, L. F. Yang, and S. R. Manalis. Mammalian cell growth dynamics in mitosis. eLife, 8:e44700, 2019.

[26] T. P. Miettinen, K. S. Ly, A. Lam, and S. R. Manalis. Single-cell monitoring of dry mass and dry mass density reveals exocytosis of cellular dry contents in mitosis. eLife, 11:e76664, 2022.

[27] K. Biswas, A. E. Sanderson, H. Salman, and N. Brenner. Single-cell growth rate variability in balanced exponential growth. bioRxiv, 2024.06.23.600237, 2024.

[28] P-Y. Ho, J. Lin, and A. Amir. Modeling cell size regulation: From single-cell-level statistics to molecular mechanisms and population-level effects. Annual Review of Biophysics, 47(1):251–271, 2018.

[29] J. Fernandez-de Cossio-Diaz, R. Mulet, and A. Vazquez. Cell population heterogeneity driven by stochastic partition and growth optimality. Sci Rep, 9(1):9406, 2019.

[30] D. Huh and J. Paulsson. Non-genetic heterogeneity from stochastic partitioning at cell division. Nat Genetics, 43(2):95–U32, 2011.

[31] G. Peruzzi, M. Miotto, R. Maggio, G. Ruocco, and G. Gosti. Asymmetric binomial statistics explains organelle partitioning variance in cancer cell proliferation. Communications Physics, 4(1):188, 2021.

[32] I. G. Johnston and N. S. Jones. Closed-form stochastic solutions for non-equilibrium dynamics and inheritance of cellular components over many cell divisions. Proc Math Phys Eng Sci, 471(2180):20150050, 2015.

[33] C. Cadart, L. Venkova, M. Piel, and M. Cosentino Lagomarsino. Volume growth in animal cells is cell cycle dependent and shows additive fluctuations. eLife, 11:e70816, 2022.

[34] L. Venkova, A. S. Vishen, S. Lembo, N. Srivastava, B. Duchamp, A. Ruppel, A. Williart, S. Vassilopoulos, A. Deslys, J. M. Garcia Arcos, A. Diz-Muñoz, M. Balland, J-F. Joanny, D. Cuvelier, P. Sens, and M. Piel. A mechano-osmotic feedback couples cell volume to the rate of cell deformation. eLife, 11:e72381, 2022.

[35] J. H. Kang, T. P. Miettinen, L. Chen, S. Olcum, G. Katsikis, P. S. Doyle, and S. R. Manalis. Noninvasive monitoring of single-cell mechanics by acoustic scattering. Nat Methods, 16(3):263–269, 2019.

[36] R. J. Kimmerling, G. Lee Szeto, J. W. Li, A. S. Genshaft, S. W. Kazer, K. R. Payer, J. de Riba Borrajo, P. C. Blainey, D. J. Irvine, A. K. Shalek, and S. R. Manalis. A microfluidic platform enabling single-cell rna-seq of multigenerational lineages. Nat Commun, 7:10220, 2016.

[37] M. Jang and H. N. Kim. From single-to multi-organ-on-a-chip system for studying metabolic diseases. BioChip Journal, 17(2):133–146, 2023.

[38] R. Kim. Advanced organotypic in vitro model systems for host–microbial coculture. Biochip journal, 17(2):147–173, 2023.

[39] A. Amir. Cell size regulation in bacteria. Physical Review Letters, 112(20):208102, 2014.

[40] M. Campos, I. V. Surovtsev, S. Kato, A. Paintdakhi, B. Beltran, S. E. Ebmeier, and C. Jacobs-Wagner. A constant size extension drives bacterial cell size homeostasis. Cell, 159(6):1433–1446, 2014.

[41] S. Taheri-Araghi, S. Bradde, J. T. Sauls, N. S. Hill, P. A. Levin, J. Paulsson, M. Vergassola, and S. Jun. Cell-size control and homeostasis in bacteria. Curr Biol, 25(3):385–391, 2015.

[42] P-Y. Ho, J. Lin, and A. Amir. Modeling cell size regulation: From single-cell-level statistics to molecular mechanisms and population-level effects. Annual Review of Biophysics, 47:251–271, 2018.

[43] M. M. Logsdon, P-Y. Ho, K. Papavinasasundaram, K. Richardson, M. Cokol, C. M. Sassetti, A. Amir, and B. B. Aldridge. A parallel adder coordinates mycobacterial cell-cycle progression and cell-size homeostasis in the context of asymmetric growth and organization. Curr Biol, 27(21):3367–3374.e7, 2017.

[44] M. Deforet, D. van Ditmarsch, and J. B. Xavier. Cell-size homeostasis and the incremental rule in a bacterial pathogen. Biophysical Journal, 109(3):521–528, 2015.

[45] L. Willis and K. C. Huang. Sizing up the bacterial cell cycle. Nat Rev Microbiol, 15:606–620, 2017.

[46] Y-J. Eun, P-Y. Ho, M Kim, S. LaRussa, L. Robert, L. D. Renner, A. Schmid, E. Garner, and A. Amir. Archaeal cells share common size control with bacteria despite noisier growth and division. Nat Microbiol, 3:148–154, 2018.

[47] Y. Sung, A. Tzur, S. Oh, W. Choi, V. Li, R. R. Dasari, Z. Yaqoob, and M. W. Kirschner. Size homeostasis in adherent cells studied by synthetic phase microscopy. Proc Natl Acad Sci U S A, 110(41):16687–92, 2013.

[48] L. Mu, J. H. Kang, S. Olcum, K. R. Payer, N. L. Calistri, R. J. Kimmerling, S. R. Manalis, and T. P. Miettinen. Mass measurements during lymphocytic leukemia cell polyploidization decouple cell cycle- and cell size-dependent growth. Proc Natl Acad Sci U S A, 117(27):15659–15665, 2020.

[49] P. Swain, K. Stevenson, A. Leary, L. F. Montano-Gutierrez, I. B. N. Clark, J. Vogel, and T. Pilizota. Inferring time derivatives including cell growth rates using gaussian processes. Nat Commun, 7:13766, 2016.

[50] P. D. Tonner, C. L. Darnell, B. E. Engelhardt, and A. K. Schmid. Detecting differential growth of microbial populations with gaussian process regression. Genome research, 27(2):320–333, 2017.

[51] C. K. I. Williams and C. E. Rasmussen. Gaussian processes for machine learning, volume 2. MIT press Cambridge, MA, 2006.

[52] X. Liu, S. Oh, L. Peshkin, and M. W. Kirschner. Computationally enhanced quantitative phase microscopy reveals autonomous oscillations in mammalian cell growth. Proc Natl Acad Sci U S A, 117(44):27388–27399, 2020.

[53] A. Amir. Thinking Probabilistically: Stochastic Processes, Disordered Systems, and Their Applications. Cambridge University Press, 2020.

[54] N. Nordholt, J. H. van Heerden, and F. J. Bruggeman. Biphasic cell-size and growth-rate homeostasis by single bacillus subtilis cells. Curr Biol, 30(12):2238–2247.e5, 2020.

[55] M. D’Ario, R. Tavares, K. Schiessl, B. Desvoyes, C. Gutierrez, M. Howard, and R. Sablowski. Cell size controlled in plants using dna content as an internal scale. Science, 372(6547):1176–1181, 2021.

[56] M. P. Swaffer, J. Kim, D. Chandler-Brown, M. Langhinrichs, G. K. Marinov, W. J. Greenleaf, A. Kundaje, K. M. Schmoller, and J. M. Skotheim. Transcriptional and chromatin-based partitioning mechanisms uncouple protein scaling from cell size. Mol Cell, 81(23):4861–4875 e7, 2021.

[57] R. Jajoo, Y. Jung, D. Huh, M. Viana, S. Rafelski, M. Springer, and J. Paulsson. Accurate concentration control of mitochondria and nucleoids. Science, 351(6269):169–172, 2016.

