## Supplementary Section for "Stochasticity in mammalian cell growth rates drives cell-to-cell variability independently of cell size and divisions"

### SI Appendix for:

[1] *Department of Mathematics, Dartmouth College, USA*
[2] *Department of Mechanical Engineering, Seoul National University, Seoul 08826, Republic of Korea*
[3] *Institute of Advanced Machines and Design, Seoul National University, Seoul 08826, Republic of Korea*
[4] *Koch Institute for Integrative Cancer Research, Massachusetts Institute of Technology, Cambridge, MA 02139, USA*
[5] *Department of Physics of Complex Systems, Weizmann Institute of Science, 7610001 Rehovot, Israel*
[6] *Department of Mechanical Engineering, Massachusetts Institute of Technology, Cambridge, MA 02139, USA*
[7] *Department of Biological Engineering, Massachusetts Institute of Technology, Cambridge, MA 02139, USA*
[8] *MRC Laboratory for Molecular Cell Biology, University College London, London, WC1E 6BT, UK*
[*] *Equal contribution*

**3 Time-dependency of cell-to-cell variability in $\lambda$** **7**

**4 Limitations of the study** **7**

**5 Supplementary Figures** **9**

### 1 Single-cell growth models

Here, we describe a simple null model for growth rates dynamics that are unregulated. This is a special case of the model studied in [1]. To begin we assume growth rate and mass dynamics within the cell-cycle are given by

$$\frac{d}{dt}\lambda(t) = \gamma(\lambda_0 - \lambda(t)) + \sqrt{2D}\xi$$
$$\frac{d}{dt}M(t) = \lambda(t)M(t).$$

At the division times $t_1 < t_2 < t_3 < \dots$ the cell size is divided symmetrically. The growth rate is perturbed at cell division and growth relaxes exponentially at a rate $\gamma$ to a lineage specific mean $\lambda_0$. If $\lambda'$ is the mother cell growth rate immediately before division and $\tilde{\lambda}$ is the growth rate of a newborn cell, then we set $z = \tilde{\lambda} - \lambda'$ and use $\sigma_z^2$ to denote the variance of the growth rate perturbation from mother to daughter. With this model we can interpolate between the limit of a pure OU process, which is blind to division events ($\sigma_z^2 = 0$), and a Division noise (DN) model, where all the noise is introduced at cell division ($D = 0$, and $\sigma_z^2 > 0$)). (see Figure S1.)

For a cell born at time $t_i$ having generation time $\tau_i$ and $t_i \le t < t_i + \tau_i = t_{i+1}$,

$$M(t) = M(t_i)e^{\int_{t_i}^{t} \lambda(s)ds}.$$

Because division is symmetric, $\lim_{t\to t_i^-} M(t) = 2M(t_i)$. The log masses obey

$$\ln M(t) = -\ln(2) + \ln M(t_i) + \int_{t_i}^{t} \lambda(s)ds \tag{1}$$
$$= -K(t)\ln 2 + \int_{0}^{t} \lambda(s)ds \tag{2}$$

where $K(t)$ is the number of cell divisions before time $t$. We will implement cell divisions by assuming that cells have the ability to "sense" their size and divide approximately at the size $f(M(t_i))$, where $f(x)$ implements the cell-size regulation strategy. We assume an adder strategy, $f(M(t_i)) = M(t_i) + \Delta$, where $\Delta$ is the size increment added between each division event.

Since the model is used as a null model for the growth process and its role in size control, the specific manner in which size control is implemented is not important. Therefore, we take the simplest approach: We assume division occurs when the mass reaches perturbation of $f(M(t_i)) + \delta_M$ where $\delta_M \sim \text{Normal}(0, \sigma_M)$. Mathematically,

$$t_{i+1} = \inf_{s>t_i}\{s : M(s) > f(M(t_i)) + \xi_i\} \tag{3}$$

where $\xi_i \sim \text{Normal}(0, \sigma_M)$ is a noise term assigned to the $i$th cell at birth.

#### 1.1 Moment dynamics

The expressions of the conditional and total mean are respectively

$$\mathbb{E}[\lambda|\tilde{\lambda}] = \tilde{\lambda}e^{-\gamma t} + \lambda_0(1 - e^{-\gamma t}) \tag{4}$$
$$\mathbb{E}[\lambda] = \mathbb{E}[\mathbb{E}[\lambda|\tilde{\lambda}]] = \lambda_0 \tag{5}$$

Similarly we obtain the conditional expression for the average of the product of $\lambda(t_1)$ and $\lambda(t_2)$ as follows

$$\mathbb{E}[\lambda_1\lambda_2|\tilde{\lambda}] = \mathbb{E}[\lambda_1|\tilde{\lambda}] \cdot \mathbb{E}[\lambda_2|\tilde{\lambda}] + \frac{D}{\gamma}\left(e^{-\gamma|t_1-t_2|} - e^{-\gamma(t_1+t_2)}\right) \quad [\lambda_1 = \lambda(t_1), \lambda_2 = \lambda(t_2)] \tag{6}$$

This expression will help to obtain the variance and covariance of $\lambda(t)$, which we will derive next. Now, if $t_1 = t_2$, then $\lambda_1 = \lambda_2$ and $\mathbb{E}[\lambda_1\lambda_2|\tilde{\lambda}] = \mathbb{E}[\lambda^2|\tilde{\lambda}]$. The conditional variance $\text{Var}(\lambda|\tilde{\lambda}) = \mathbb{E}[\lambda^2|\tilde{\lambda}] - (\mathbb{E}[\lambda|\tilde{\lambda}])^2$ can be calculated from Eq. (6), which is given as follows

$$\text{Var}(\lambda|\tilde{\lambda}) = \frac{D}{\gamma}\left(1 - e^{2\gamma t}\right). \tag{7}$$

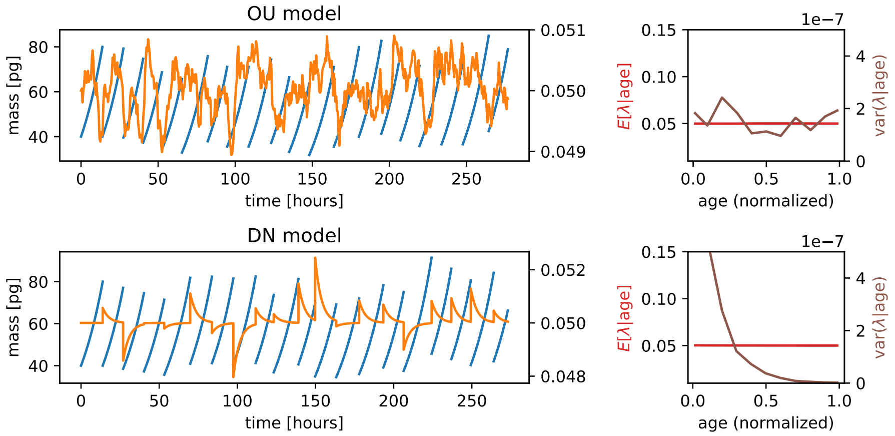

Figure S1: (top) Simulations of OU model showing mass (blue) and growth rate (orange) dynamics along with the variance and expected value of growth rates conditioned on age (top right). This illustrates that both are nearly independent of age. (bottom) The same figure for the Division noise model.

Using the law of total variance we have

$$\begin{aligned}\mathrm{Var}(\lambda) &= \mathrm{Var}(\mathbb{E}[\lambda|\tilde{\lambda}]) + \mathbb{E}[\mathrm{Var}(\lambda|\tilde{\lambda})] \\ &= \mathrm{Var}(\tilde{\lambda}e^{-\gamma t} + \lambda_0(1-e^{-\gamma t})) + \mathbb{E}\left[\frac{D}{\gamma}\left(1-e^{-2\gamma t}\right)\right] \\ &= \mathrm{Var}(\tilde{\lambda})e^{-2\gamma t} + \frac{D}{\gamma}\left(1-e^{-2\gamma t}\right) \end{aligned} \tag{8}$$

Now for our problem $\tilde{\lambda}$ is actually OU process at steady-state, implying $\mathbb{E}[\tilde{\lambda}] = \lambda_0$ and $\mathrm{Var}(\tilde{\lambda}) = D/\gamma$. But due to uneven division there will be another noise term in $\tilde{\lambda}$. We denote the variance in $\tilde{\lambda}$ due to division is $\sigma_z^2$. So $\mathrm{Var}(\tilde{\lambda}) = (D/\gamma + \sigma_z^2)$. Now using the expression of $\mathrm{Var}(\tilde{\lambda})$ in Eq. (8) we obtain

$$\mathrm{Var}(\lambda) \approx \frac{D}{\gamma} + \sigma_z^2 e^{-2\gamma t}. \tag{9}$$

Similarly, using the law of total covariance we obtain

$$\mathrm{Cov}(\lambda(t_1), \lambda(t_2)) = \frac{D}{\gamma}e^{-\gamma|t_1-t_2|} + \sigma_z^2 e^{-\gamma(t_1+t_2)}. \tag{10}$$

Now, the variance of the time averaged growth rate $\lambda_t = \int_0^t \lambda(s)ds$ will be

$$\begin{aligned}\mathrm{Var}(\bar{\lambda}) &= \frac{1}{\tau^2}\int_0^\tau\int_0^\tau \mathrm{Cov}(\lambda(t_1),\lambda(t_2))dt_1dt_2 & (11)\\ &= \frac{2D}{\gamma^2\tau^2}\left[\tau - \frac{1}{\gamma}(1-e^{-\gamma\tau})\right] + \left(\frac{\sigma_z}{\gamma\tau}\right)^2(1-e^{-\gamma\tau}). & (12)\end{aligned}$$

When $\gamma\tau > 1$ this expression will reduce into the following form

$$\mathrm{Var}(\bar{\lambda}) = \frac{1}{\gamma^2}\left(\frac{2D}{\tau} + \frac{\sigma_z^2}{\tau^2}\right). \tag{13}$$

#### 1.2 Model parameters used in simulated datasets

In are simulated datasets used to make Fig 5 in the main text, we fixed the average growth rate and the average cell-size and variance in birth size ($\sigma_M^2$) according to the values in Table 1.

| Parameter | Mean | Standard Deviation | Units |
|---|---|---|---|
| $\Delta$ | 35.45 | 2.27 | Mass [pg] |
| $\lambda_0$ | 0.074 | 0.011 | 1/hours |
| $\sigma_M$ | 16.5 | 7.09 | pg |
| $\sigma_{\bar{\lambda}}$ | 0.014 | 0.003 | 1/hours |

Table 1: Table of parameter mean and standard deviations over lineages

### 2 Gaussian process smoothing and decomposition

This section explains how the dynamics of the growth rates over the course of a lineage are inferred using Gaussian process decomposition.

#### 2.1 Setup

It is useful to define the continuous function

$$f(t) = \ln \frac{M_i(t)}{M_i^b} + \sum_{j<i} \ln \frac{M_j^d}{M_j^b} = \int_0^t \lambda(s) ds \tag{14}$$

where $M_i^b$ and $M_i^d$ are the mass of the $i$th cell as birth and division.

The basic assumption of our analysis is that the growth rate can be decomposed into contributions from various factors, e.g., coupling to cell cycle progression, as

$$\lambda(t) = \lambda_0 + \sum_k \lambda_k(t) \tag{15}$$

where each term $g_k(t)$ represents the contribution from some factor. For example, for the contribution from purely age dependent effects, we would have

$$g_k(t) = h(\text{age}(t)) \tag{16}$$

where age$(t)$ converts the time since the beginning of the lineage to the (normalized) age of a cell; that is,

$$\text{age}(t) = \sum_j \mathbf{1}_{t_j<t<t_{j+1}} \frac{t - t_j}{t_{j+1} - t_j} \tag{17}$$

where $t_j$ is the division time when the $j$ cell divides. Terms depending on unnormalized age or cell-mass could be defined in a similar manner. The decomposition of $\lambda$ induces a decomposition of $f(t)$:

$$f(t) = \lambda_0 t + \sum_{k=0} f_k(t). \tag{18}$$

where $f_k(t) = \int_0^t \lambda_k(s) ds$.

#### 2.2 Mathematics of Gaussian process decomposition

A process $X_t$ is said to be a *Gaussian process* if for any finite set of observations, $\mathbf{t} = (t_1, \ldots, t_k)^T$, the joint distribution of $\mathbf{X}(\mathbf{t}) = (X_{t_1}, \ldots, X_{t_k})^T$ is a multivariate Gaussian [2]. We will typically work with zero mean Gaussian processes, so that the distribution of $X_t$ is specified entirely by a covariance function, or *kernel*:

$$k(t, t') = \mathbb{E}[X_t X_{t'}]. \tag{19}$$

We will use $K(\mathbf{t})$ to represent the matrix with entries

$$K(\mathbf{t})_{i,j} = k(t_i, t_j). \tag{20}$$

Then, the distribution of a Gaussian process with covariance kernel $k$ at sample points $\mathbf{t}$ is

$$P_{\text{gp}}(\mathbf{x}; k) = \frac{1}{(2\pi)^k \det K(\mathbf{t})^{1/2}} e^{-\mathbf{x}^T K(\mathbf{t})^{-1}\mathbf{x}/2}. \tag{21}$$

Typically one selects the kernel from some parametric family, for example, a common choice is the squared exponential (SE) kernel

$$k(t, t') = Ae^{-(t-t')^2/\tau}. \tag{22}$$

Samples of a Gaussian process with an SE kernel will be smooth, however, with a different choice of kernel we can easily generate functions with different properties. We will discuss the selection of kernels in more detail later.

In a Gaussian process regression, one places Gaussian process priors on an unknown function. In the present context, this means placing a Gaussian process priors on each of the functions $f_i$ in Equation 18. In addition, we must account for the linear term. For the moment, we will ignore how the kernels and their corresponding parameters are selected and simply assume we have some kernel $k_i$ for each term $f_i$ in the decomposition. The Bayesian posterior of the vectors $\mathbf{f}_i = (f_i(t_1), \dots, f_i(t_j))^T$ is given by

$$P(\mathbf{f}_1, \dots, \mathbf{f}_L|\mathbf{y}) \propto P(\mathbf{y}|\mathbf{f}_1, \cdots, \mathbf{f}_L) \prod_i P_{\text{gp}}(\mathbf{f}_i; k_i). \tag{23}$$

Under the assumption that measurement errors are Gaussian and uncorrelated, $P(\mathbf{y}|\mathbf{f}_1, \cdots, \mathbf{f}_L)$ is a multivariate normal distribution with mean $\sum \mathbf{f}_j$ and covariance matrix $I\sigma_\epsilon^2$. The log posterior is therefore given by

$$-\ln P(\mathbf{f}_1, \dots, \mathbf{f}_L|\mathbf{y}) \propto \frac{1}{2\sigma_\epsilon^2}\left(\mathbf{y} - \sum_j \mathbf{f}_j\right)^T I \left(\mathbf{y} - \sum_j \mathbf{f}_j\right) + \frac{1}{2}\sum_j \mathbf{f}_i^T K_j(\mathbf{t})^{-1}\mathbf{f}_i \tag{24}$$

Here $\sigma_\epsilon^2$ the variance of the measurement errors. This implies that under the Gaussian process model, the marginal distribution of $\mathbf{y}$ is Gaussian mean zero with covariance matrix

$$M(\mathbf{t}) = \sum_j K_j(\mathbf{t}) + \sigma_\epsilon^2 I. \tag{25}$$

It follows that the joint distribution of $(\mathbf{y}, \mathbf{f}_i, \mathbf{f}_j)$, the data and $i$th and $j$th terms in the decomposition, is Gaussian with mean zero [1] and covariance matrix

$$\bar{K}_{i,j}(\mathbf{t}) = \begin{bmatrix} \sum_m K_m(\mathbf{t}) + I/\sigma_\epsilon^2 & K_i(\mathbf{t}) & K_j(\mathbf{t}) \\ K_i(\mathbf{t}) & K_i(\mathbf{t}) & \mathbf{0} \\ K_j(\mathbf{t}) & \mathbf{0} & K_j(\mathbf{t}) \end{bmatrix} \tag{26}$$

where $\mathbf{0}$ is a matrix of all zeros with the appropriate dimensions.

Using standard identities for the conditional mean and variance of a multivariate Gaussian, we can easily obtain the relevant statistics of the posterior. In particular, the maximum likelihood estimate of $\mathbf{f}_k$ under the posterior distribution is

$$\boldsymbol{\mu}_j(\mathbf{y}) \equiv \mathbb{E}[\mathbf{f}_j|\mathbf{y}] = K_j(\mathbf{t})M(\mathbf{t})^{-1}\mathbf{y}. \tag{27}$$

This represents our best estimate of the unknown function $\mathbf{f}_k$ given the data. Note that Equation 27 has the form of a linear smoother of the data. Also of interest is variance of $f_j(t)$ and the covariance between $f_j(t)$ and $f_i(t)$ under the posterior. The former quantifies the uncertainty in our inference while the later tells us how “sloppy” our model is, that is, whether it is difficult to distinguish between terms in our decomposition. In order to obtain the covariance, we consider the joint distribution of $f_j(t)$ and $f_i(t)$ conditional on $\mathbf{y}$, which has mean $(\boldsymbol{\mu}_i(\mathbf{y}), \boldsymbol{\mu}_j(\mathbf{y}))$ and covariance matrix

$$\begin{aligned} \bar{K}_{i,j}(\mathbf{t}|\mathbf{y}) &= \begin{bmatrix} K_i(\mathbf{t}) & \mathbf{0} \\ \mathbf{0} & K_j(\mathbf{t}) \end{bmatrix} - \begin{bmatrix} K_i(\mathbf{t}) \\ K_j(\mathbf{t}) \end{bmatrix} M(\mathbf{t})^{-1} \begin{bmatrix} K_i(\mathbf{t}) & K_j(\mathbf{t}) \end{bmatrix} \\ &= \begin{bmatrix} K_i(\mathbf{t}) - K_i(\mathbf{t})M(\mathbf{t})^{-1}K_i(\mathbf{t}) & K_i(\mathbf{t})M(\mathbf{t})^{-1}K_j(\mathbf{t}) \\ K_j(\mathbf{t})M(\mathbf{t})^{-1}K_i(\mathbf{t}) & K_j(\mathbf{t}) - K_j(\mathbf{t})M(\mathbf{t})^{-1}K_j(\mathbf{t}) \end{bmatrix}. \end{aligned} \tag{28}$$

[1] We can compute the full joint distribution of the entire decomposition, but since this is Gaussian it is sufficient to determine he distribution for any pair.

From this matrix we obtain the covariances under the posterior

$$\mathbf{C}_{i,j}(\mathbf{y}) = \begin{bmatrix} \mathrm{cov}(f_i(t_1), f_j(t_1)|\mathbf{y}) \\ \vdots \\ \mathrm{cov}(f_i(t_k), f_j(t_k)|\mathbf{y}) \end{bmatrix} = \mathrm{diag}\left(K_i(\mathbf{t})M(\mathbf{t})^{-1}K_j(\mathbf{t})\right) \tag{29}$$

Similarly, the variance is given given by

$$\boldsymbol{\nu}_j(\mathbf{y}) \equiv \mathrm{var}(\mathbf{f}_j|\mathbf{y}) = \mathrm{diag}\left(K_j(\mathbf{t}) - K_j(\mathbf{t})M(\mathbf{t})^{-1}K_j(\mathbf{t})\right). \tag{30}$$

#### 2.3 Model specifics

As discussed in the main text, our model is of the form

$$f(t) = \lambda_0 t + f_{\mathrm{trend}}(t) + f_{\mathrm{flucs}}(t) + \epsilon(t). \tag{31}$$

where $f_{\mathrm{trend}}(t)$ is a smooth function depending only on the cell's normalized age $a(t) = (t - t_i)/(t_{i+1} - t_i)$. This was achieved with a smooth kernel depending only on $|a(t) - a(t')|$.

$$k_{\mathrm{trend}}(t, t') = A_{\mathrm{trend}} e^{-|a(t)-a(t')|^2/\gamma_{\mathrm{trend}}}$$

We found that using un-normalized age makes little difference in our results, since generation time variability is relatively small. For the term $f_{\mathrm{flucs}}(t)$, we used a once differentiable kernel known as the Matérn-32 kernel depending on the distance $|t - t'|$ [2]:

$$k_{\mathrm{flucs}}(t, t') = A_{\mathrm{flucs}} \left(1 + \frac{|t - t'|}{\gamma_{\mathrm{flucs}}}\right) e^{-|t-t'|^2/\gamma_{\mathrm{flucs}}}$$

The Matérn-32 kernel corresponds to the critical regime of a stochastic, damped harmonic oscillator and thus is desirable as a prior distribution because it can accommodate both over and underdamped fluctuations.

Both kernels are parameterized by a prefactor and a time scale, which were inferred along with the noise magnitude by maximizing the marginal likelihood (collectively, we refer to these as hyper-parameters). The posterior distribution of $\lambda(t) = f'(t)$ with fixed values of the hyperparameters can then be obtained analytically.

#### 2.4 Model selection and kernel parameter optimization

To evaluate the fit of our model, we use the marginal log likelihood $P(\mathbf{y})$, which is the distribution of our data, marginalized over all the Gaussian process priors. This will depend on both the model (i.e. the number of terms in our decomposition and the kernels we have selected for them), as well as the kernel parameters. Thus, we write

$$P(\mathbf{y}) = P\left(\mathbf{y}|\{k_j, \boldsymbol{\theta}_j\}_{j=1}^L\right) \tag{32}$$

where $k_j$ are the kernels and $\boldsymbol{\theta}_j$ are their parameters. Together these uniquely define the priors. By performing a multivariate Gaussian integral, it can be shown that

$$\ln P\left(\mathbf{y}|\{k_j, \boldsymbol{\theta}_j\}_{j=1}^L\right) = -\frac{1}{2}\mathbf{y}^T M(\mathbf{t})^{-1}\mathbf{y} - \frac{1}{2}\ln|M(\mathbf{t})| - \frac{k}{2}\ln 2\pi \tag{33}$$

By maximizing $\ln P(\mathbf{y}|\{k_j, \boldsymbol{\theta}_j\}_{j=1}^L)$ with respect to the $\boldsymbol{\theta}_j$'s, we can obtain the most likely parameters for our kernels, which are then used in our inference of the unknown functions $G_j$. Note that in a fully Bayesian approach we would place prior distributions on these parameters and obtain their posterior distribution as well. However, the posterior distribution of the kernel parameters will generally be non-Gaussian and analytically intractable, thus we would need to utilize Markov chain Monte Carlo simulations which are computationally expensive. Moreover, with the present quantity of data it is sufficient to treat these parameters as fixed, since the posterior distribution of $G_j$ will generally be insensitive to their values within a small neighborhood.

### 3 Time-dependency of cell-to-cell variability in $\lambda$

Some cellular perturbations are considered to influence fast growing cells differently from slow growing cells. In bacteria, fast growing cells are more likely to die following certain antibiotic treatments [3], and in cancer cells, slow growing cells are often more resilient to chemotherapy [4, 5, 6]. As many drugs are rapidly cleared from the plasma [7], there is a narrow time window during which such treatments are effective. Cell-to-cell variability in $\lambda$ could therefore influence the fraction of cells affected by the drug. Yet, the magnitude of cell-to-cell variability in $\lambda$ is difficult to assess from the literature, as single-cell growth studies are carried out with different time-resolutions. To address this in our data, we randomly selected time points from each lineage and compared the cell-to-cell variability in growth rates when $\lambda$ was analyzed using different averaging times. The growth rates on 1 minute timescale contained $\sim$15% cell-to-cell variability within each lineage, and this gradually decreased to $\sim$7% as the growth rate timescale approached the typical interphase duration (i.e. $\sim \bar{\lambda}$, Figure S2). Thus, the variability in leukemia cell growth rates is $\sim$100% higher on short timescales ($\leq$0.5 hour) than on long timescales ($\geq$6 hours). Consequently, a short exposure to a drug which targets only fast growing cells, is likely to impact a significantly smaller fraction of cells than a long drug exposure simply due to the fluctuating nature of $\lambda$. By revealing the time-dependency of cell-to-cell variability in $\lambda$, our work provides an approach for optimizing drug exposure times when the goal is to minimize the heterogeneity in cellular responses to the drug.

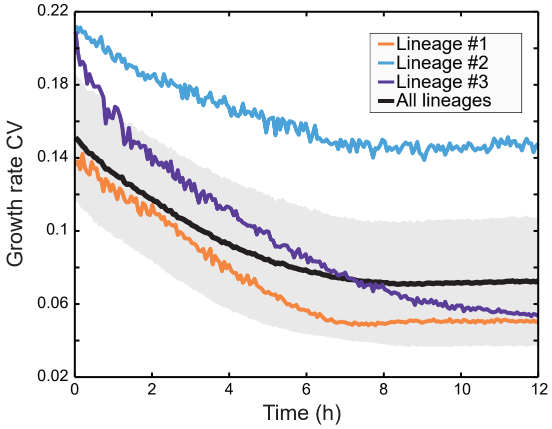

Figure S2: Dependency of cell-to-cell $\lambda$ variability on data averaging time. Data is displayed for the three longest lineages (colored lines). The black line and grey shaded area represent lineage mean±SD ($N = 24$ independent lineages/experiments).

### 4 Limitations of the study

A key obstacle in the study of fluctuations in $\lambda$ is the separation of environmental and measurement fluctuations from cell-intrinsic growth fluctuations. While we have excluded measurement fluctuations by comparing our cell measurements to polystyrene bead measurements, we cannot fully exclude environmental fluctuations that could influence cell growth rates. However, as our data is gathered from multiple measurement setups over several years, environmental fluctuations are unlikely to have a major impact on the average cell behavior we report. However, a potential exception to this is the cell lineage #2, where the GP analysis was not able to fully capture the fine details of the fluctuations in $\lambda$, as evident by the higher correlation between consecutive residuals (Fig. S5B). The poor performance of the GP analysis regarding this lineage could reflect environmental changes.

Another important limitation of our work is the fact that our biological conclusions are limited to a single cell line and a single growth environment. It is reasonable to assume that the magnitude of growth noise in a given model system depends on the genetic background of the cells, as well as the nutrients and growth signaling that the cell is receiving.

Our work also shows that each cell lineage has its typical growth rate (Fig. 1B), but the stochastic cell-to-cell variability we identify within lineages does not explain this lineage-to-lineage variability. Our data did not reveal clear instances where ancestral lineages grow at their typical rate and then shift to another steady growth rate. We speculate that observing such transitions in the lineage-specific growth rate could require monitoring of growth over significantly longer lineages (>100 generations), especially if the lineage-specific growth rates are set by genetic drift.

### References

[1] Y. Hein and F. Jafarpour. Asymptotic decoupling of population growth rate and cell size distribution. *Phys. Rev. Res.*, 6(4):043006, 2024.

[2] C. K. I. Williams and C. E. Rasmussen. *Gaussian processes for machine learning*, volume 2. MIT press Cambridge, MA, 2006.

[3] N. M. V. Sampaio, C. M. Blassick, V. Andreani, J-B. Lugagne, and M. J. Dunlop. Dynamic gene expression and growth underlie cell-to-cell heterogeneity in escherichia coli stress response. *Proc Natl Acad Sci U S A*, 119(14):e2115032119, 2022.

[4] S. Shen, S. Vagner, and C. Robert. Persistent cancer cells: The deadly survivors. *Cell*, 183(4):860–874, 2020.

[5] C. Duy, M. Li, M. Teater, C. Meydan, F. E. Garrett-Bakelman, T. C. Lee, C. R. Chin, C. Durmaz, K. C. Kawabata, E. Dhimolea, C. S. Mitsiades, H. Doehner, R. J. D'Andrea, M. W. Becker, E. M. Paietta, C. E. Mason, M. Carroll, and A. M. Melnick. Chemotherapy induces senescence-like resilient cells capable of initiating aml recurrence. *Cancer Discovery*, 11(6):1542–1561, 2021.

[6] K. A. Fennell, D. Vassiliadis, E. Y. N. Lam, L. G. Martelotto, J. J. Balic, S. Hollizeck, T. S. Weber, T. Semple, Q. Wang, D. C. Miles, L. MacPherson, Y. C. Chan, A. A. Guirguis, L. M. Kats, E. S. Wong, S. J. Dawson, S. H. Naik, and M. A. Dawson. Non-genetic determinants of malignant clonal fitness at single-cell resolution. *Nature*, 601(7891):125–131, 2022.

[7] D. R. Liston and M. Davis. Clinically relevant concentrations of anticancer drugs: A guide for nonclinical studies. *Clinical Cancer Research*, 23(14):3489–3498, 2017.

[8] R. J. Kimmerling, G. Lee Szeto, J. W. Li, A. S. Genshaft, S. W. Kazer, K. R. Payer, J. de Riba Borrajo, P. C. Blainey, D. J. Irvine, A. K. Shalek, and S. R. Manalis. A microfluidic platform enabling single-cell rna-seq of multigenerational lineages. *Nat Commun*, 7:10220, 2016.

### 5 Supplementary Figures

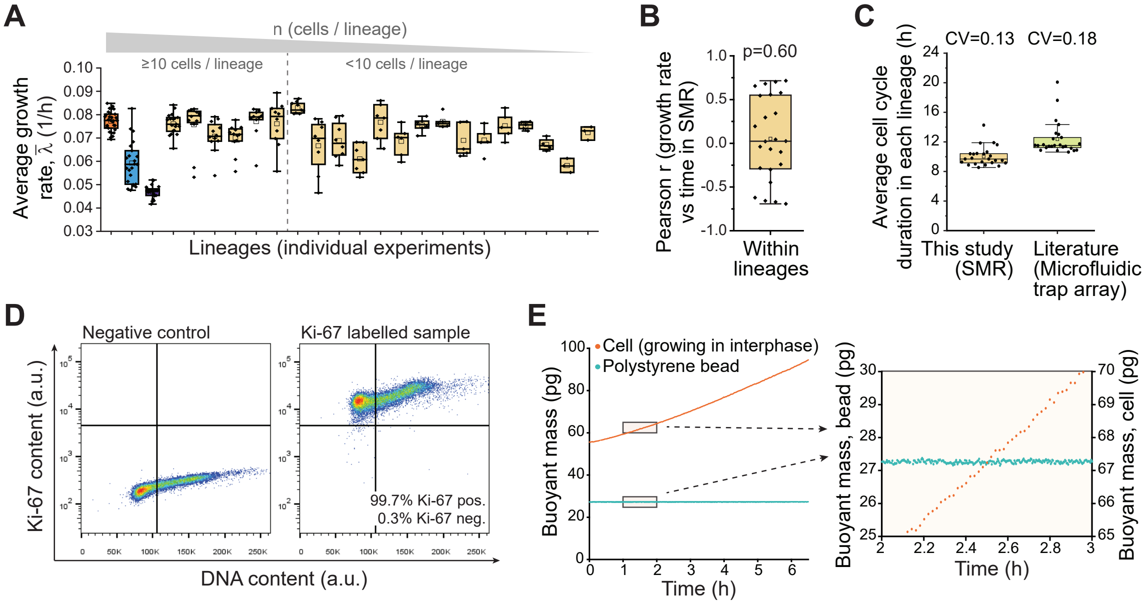

Figure S3: High-precision monitoring of leukemia cell growth under steady, high-growth conditions (A) Average cell growth rates ($\bar{\lambda}$) across all data. Each box represents an independent lineage (independent experiment, N=24) and each dot represents the $\bar{\lambda}$ of a cell in the lineage. (B) Pearson correlation value (r) for average cell growth rates ($\bar{\lambda}$) in the SMR as a function of time that the experiment had lasted. N=24 independent lineages/experiments. (C) Average cell cycle duration ($\tau_d$) within a lineage, as observed in the SMR (N=24) and using an independent microfluidic trap array (N=25) (data from [8]). The SMR data is collected with only one lineage per experiment, while the microfluidic trap array measures multiple lineages per experiment. Coefficient of variability of $\tau_d$ values is shown on top. (D) Representative Ki-67 and DNA labeling in L1210 cells. Over 99.5% of L1210 cells in a population are Ki-67 positive, indicating that virtually all cells are actively proliferating (N=3 independent experiments). (E) Representative buoyant mass monitoring data for a cell (red) and a polystyrene bead (teal, representative of 5 independent bead experiments). Inset on right displays a zoom-in.

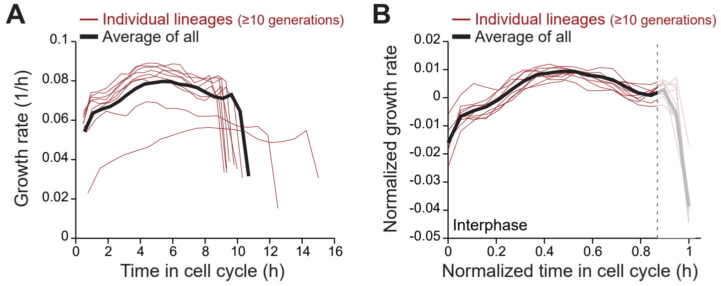

Figure S4: Cell-cycle-dependent growth corresponds to cell age (normalized time in the cell cycle) (**A**) Cell growth rates across the cell cycle as a function of time since birth. Orange lines are averages within individual lineages. Black line is the average of all shown lineages. Decreased growth rates at the end of the cell cycle is due to mitosis, which is excluded from our other analyses. (**B**) Same as (A), but data are plotted as a function of cell age (normalized time in the cell cycle). Only lineages with at least 10 full cell cycles were included (N=9 independent lineages/experiments).

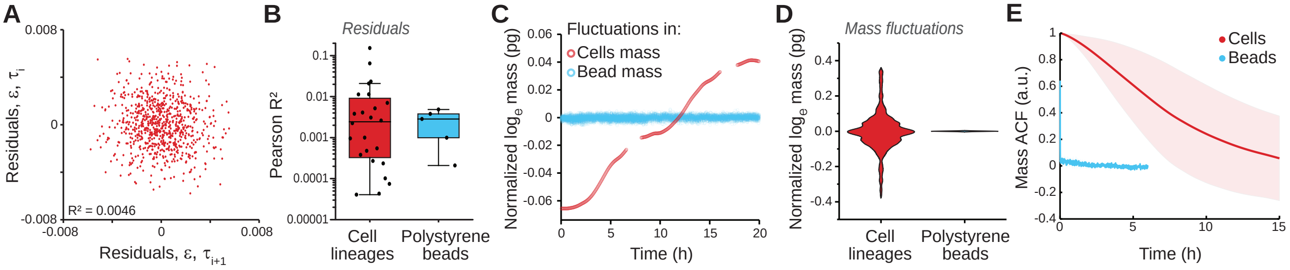

Figure S5: Gaussian process analysis captures biological mass fluctuations (**A**) A representative example of the correlation between consecutive residuals of the Gaussian progress regression analysis within one lineage. (**B**) Pearson correlation $R^2$ values for consecutive residuals in all cell lineages. Correlations are similar in magnitude to what would be expected from pure measurement noise, as shown by comparison to correlations between consecutive measurements of single polystyrene bead's mass. The cell lineage with the highest correlation is lineage #2. Each point is an independent lineage/experiment. (**C**) Representative example of GP isolated cell mass fluctuations (term $f_3(t)$, red) together with mass fluctuation observed in a polystyrene bead (blue). (**D**) Comparison between mean-normalized GP isolated cell mass fluctuations (term $f_3(t)$) in all lineages (red) and mean-normalized mass fluctuations observed in polystyrene bead measurements (blue). The mass fluctuations in cells are significantly larger than in polystyrene beads, indicating that the mass fluctuations in cells are not measurement artefacts. (**E**) Autocorrelation functions for GP isolated cell mass fluctuations (term $f_3(t)$) (red) and for polystyrene bead mass fluctuations (blue). Thick lines and shaded areas represent mean $\pm$ SD. N=24 independent lineages/experiments for all cell data. N=5 independent experiments for all bead data.

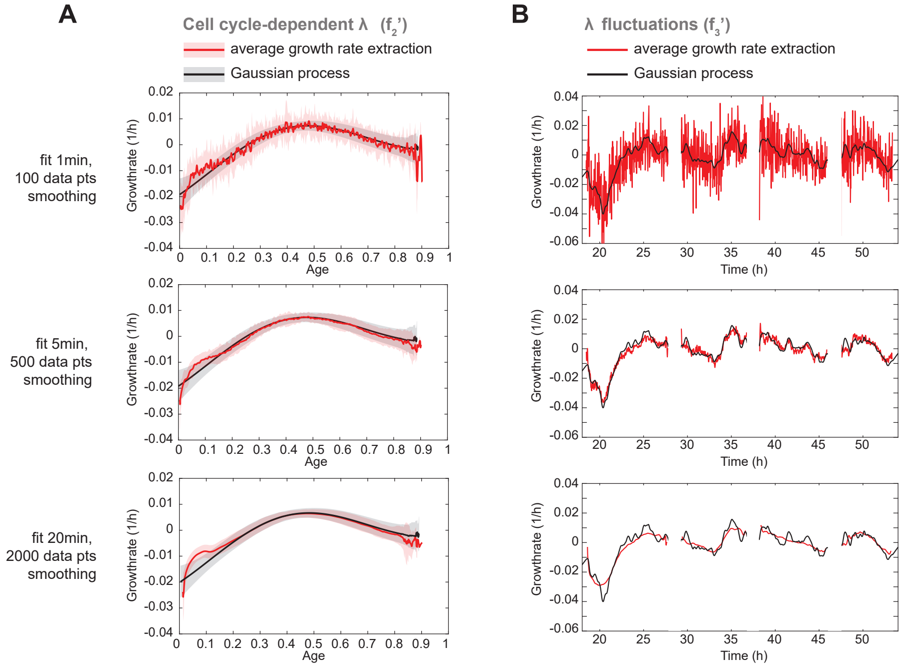

Figure S6: Growth rate comparison using GP and average growth rate extraction. (**A**) Cell cycle-dependent growth rates were extracted using GP or 'naive' data fitting with three different levels of data smoothing and fitting length. Thick lines and shaded areas represent mean $\pm$ SD. (**B**) The $\lambda$ fluctuations were extracted using GP or 'naive' data fitting with three different levels of data smoothing and fitting length.

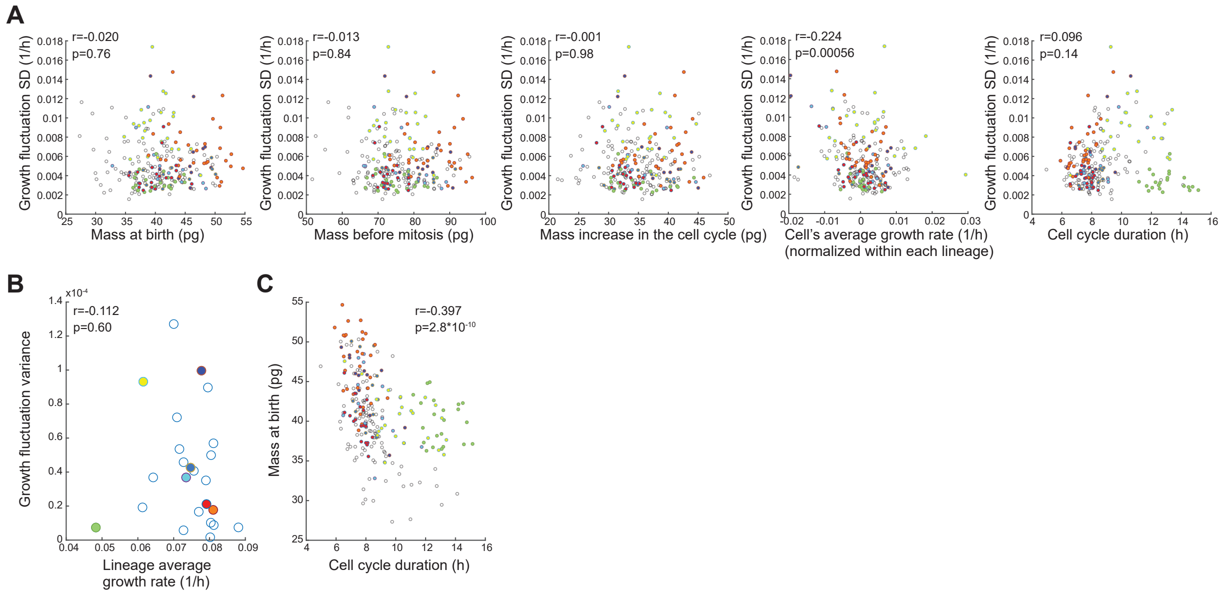

Figure S7: The $\lambda$ fluctuations are not explained by cell size-dependent growth. (**A**) Correlations between the magnitude of the $\lambda$ fluctuations (standard deviation of the $\lambda$ fluctuations across a cell cycle) and indicated cell mass and growth parameters. Each dot represents a single cell. Cells from the longest lineages are color coded. (**B**) Correlations between the magnitude of the $\lambda$ fluctuations (variance of the fluctuation term across a lineage) and the average growth rate of each lineage. Each dot represents a lineage. Longest lineages are color coded as in panel (A). (**C**) Correlations between cell mass at birth and cell cycle duration. Each dot represents a single cell. Cells from the longest lineages are color coded. In all panels, N=24 independent lineages/experiments with 235 cells in total.
